# Endosome motility controls light-responsive reproductive development and secondary metabolite production in *Aspergillus*

**DOI:** 10.64898/2026.03.03.708097

**Authors:** Gaurav Kumar, Jessica L. Allen, Livia D. S. Oster, Mira Syahfriena Amir Rawa, Enrique A. Ramírez, Jin Woo Bok, Patreece H. Suen, Bellana E. Driscoll, John Salogiannis, Nancy P. Keller, Samara L. Reck-Peterson

**Author notes:** Correspondence should be addressed to S.L.R-P. These authors contributed equally (Gaurav Kumar, Jessica L. Allen, Livia D. S. Oster).

## Abstract

Filamentous fungi, such as *Aspergillus* species, use microtubule transport to move early endosomes. Other cargos, such as peroxisomes and mRNAs, “hitchhike” on early endosomes to move throughout the long hyphae of these organisms. In *Aspergillus nidulans*, peroxisomes hitchhike on early endosomes using the endosomal protein PxdA and the peroxisomal protein AcbdA. The HookA adaptor protein links endosomes to microtubule motors. Here, we set out to explore the physiological functions of peroxisome hitchhiking and endosome motility. *Aspergillus nidulans* has a complex life cycle that includes asexual and sexual reproduction. *A. nidulans* and other fungi within the Pezizomycotina subphylum are also notable for the vast number of secondary metabolites they produce. Light and other environmental conditions influence developmental decisions and secondary metabolite production. Here, we found that sexual reproduction is favored in the absence of endosome motility, even in the light, which normally promotes asexual reproduction. RNA sequencing of strains lacking PxdA-marked motile early endosomes showed altered expression of genes involved in development. Unexpectedly, we also observed altered expression of genes involved in secondary metabolism in strains lacking endosome motility and peroxisome hitchhiking. Using mass spectrometry, we found that the loss of endosome motility affected the biosynthesis of secondary metabolites, including sterigmatocystin, a carcinogenic mycotoxin that is a food contaminant. Finally, in a pathogenic species, *Aspergillus fumigatus*, we found that deletion of its *pxdA* homolog also significantly altered secondary metabolite production. Our work uncovers an unexpected link between organelle motility, developmental decisions in response to light, and secondary metabolite production in filamentous fungi.

## Introduction

In eukaryotic cells, most large intracellular objects are moved and positioned by active transport along the cytoskeleton. In both metazoan cells and filamentous fungi, long distance cargo transport is accomplished by dynein and kinesin microtubule motors. Typically both types of motors use adaptor proteins to link directly to their cargos^1–7^. In contrast to this canonical mode of transport, some cargos do not directly recruit the transport machinery, and instead they move by attaching to other motile cargos^1,4,8^. This type of non-canonical transport termed ‘hitchhiking’ was originally described in filamentous fungi for the hitchhiking of mRNAs^9,10^ and organelles^11,12^. It is now appreciated that hitchhiking is used across the tree of life^4^.

Organelle hitchhiking was first described in two species of filamentous fungi: *Aspergillus nidulans*^11^ and the plant pathogen *Ustilago maydis*^12^. From a genetic screen designed to identify genes required for nuclei, endosome, or peroxisome positioning^13^, we identified the gene *pxdA* as being required for peroxisome localization but not nuclei or endosome distribution in *A. nidulans*^11^. Live-cell imaging in different mutant backgrounds revealed that PxdA was an endosomal protein required for peroxisomes to attach to moving early endosomes^11^. Later, the endosomal phosphatase DipA was shown to be required for peroxisome hitchhiking^14^ and the acyl-CoA binding protein, AcbdA, was identified as the first peroxisome-localized protein required for peroxisome hitchhiking^15^. Endosomes in *A. nidulans*, as is the case in metazoan cells, use a Hook adaptor protein (HookA in *A. nidulans*) to connect endosomes to microtubule motors^16^. Our current model of endosome motility and organelle hitchhiking is shown in Figure 1A.

**Figure 1.**
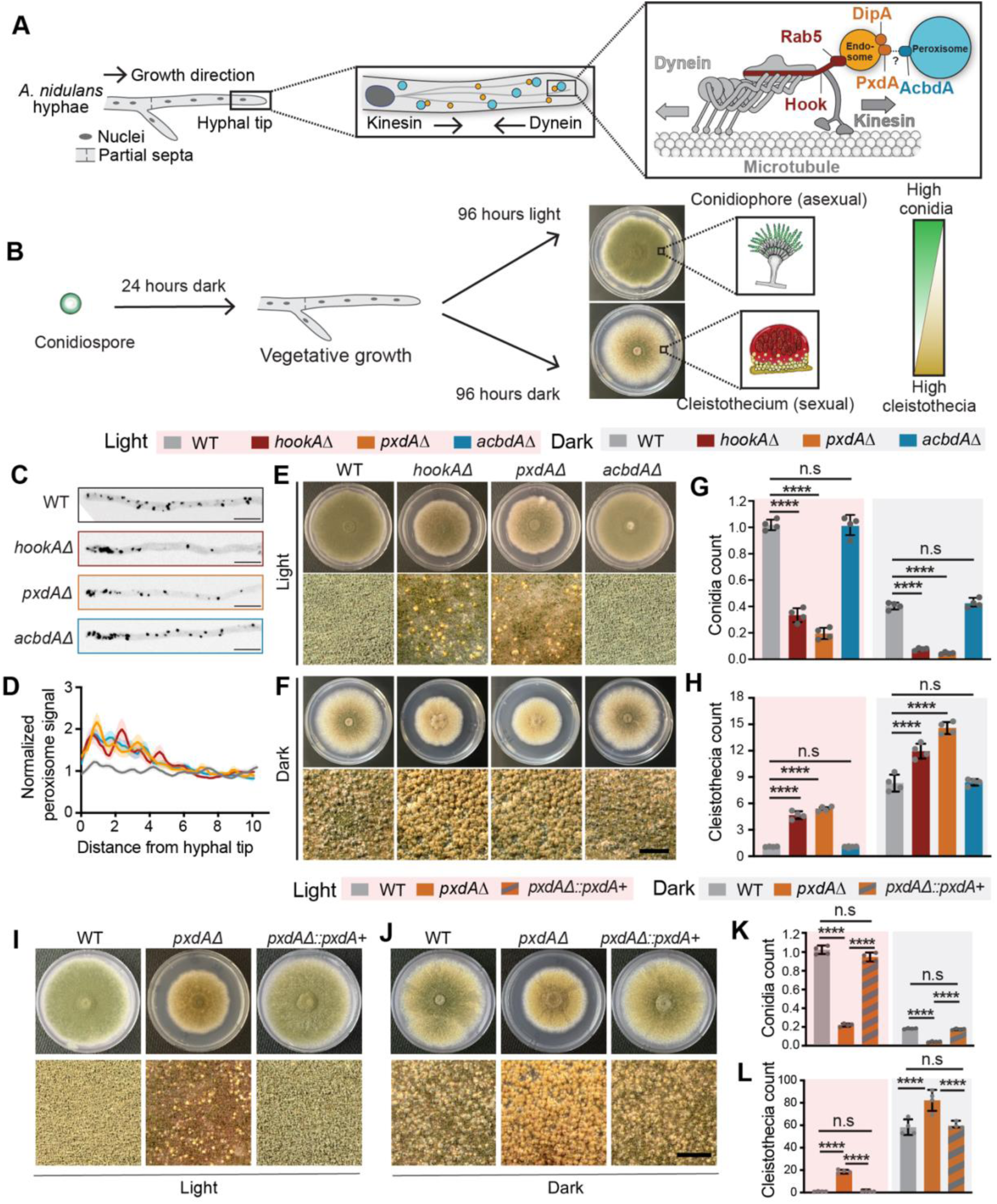
Endosomal proteins are required for light-responsive fungal development. **(A**) Schematic of vegetatively growing hyphal-tip and model of long-distance endosome motility and peroxisome hitchhiking in A. nidulans hyphae (right). Peroxisome hitchhiking on early endosomes requires the endosomal proteins PxdA and DipA and the peroxisomal protein AcbdA. HookA recruits dynein and kinesin to endosomes for microtubule motility. **(B)** Developmental morphology of A. nidulans under light versus dark conditions. The middle panels show representative colony images; the inset schematic depicts the dominant reproductive structures (asexual conidiophores in light, sexual cleistothecia in darkness). The adjacent color bar illustrates the shift from high conidiation (green) under light to high cleistothecia formation (yellow) in the dark. (**C**) Representative micrographs of wild-type, hookAΔ, pxdAΔ, and acbdAΔ hyphae showing the localization of peroxisomes labeled with GFP-AcuE. (**D**) Graph of a line scan analysis for peroxisome distribution. The line scan runs from the hyphal tip to the first nucleus, n >70 hyphae per strain. (**E** and **F**) Image of agar plate showing growth phenotype for wild-type, hookAΔ, pxdAΔ, and acbdAΔ strains grown in light (**E**) and dark (**F**). (**G**) Quantification of conidia normalized to the wild-type strain grown in light. (**H**) Quantification of cleistothecia normalized to the wild-type strain grown in light. (**I** and **J**) Image of agar plate showing growth phenotype of wild-type, pxdAΔ, and pxdAΔ::pxdA+ strains grown on glucose minimal media under light (**I)** and dark (**J**) condition. The repression of cleistothecia production in presence of light was restored by complementation of pxdA+ allele (**K**) Quantification of conidia normalized to the wild-type strain grown in light. (L) Quantification of cleistothecia normalized to the wild-type strain grown in light. Data are presented as mean ± SD (n = 4 biological replicates). **** p < 0.0001, n.s (not significant) by two-way ANOVA. Scale bars: 10 µm in (C) 2mm in (E, F, I and J) .

The *pxdA* gene is found specifically within the Pezizomycotina subphylum of fungi^17^, whereas *dipA*^17^ and *acbdA*^15^ are more widely distributed. The Pezizomycotina contain over 85,000 known fungal species^18^, encompassing ecologically essential and directly human-relevant biodiversity. Nearly 50% of the most concerning human fungal pathogens as determined by the World Health Organization are in the Pezizomycotina^19^, as are most plant pathogenic species^20^. Among its members is *Aspergillus fumigatus*, one of the most pathogenic fungal species in humans, responsible for life-threatening diseases such as invasive and chronic pulmonary aspergillosis^21,22^.

The metabolic capacity of the Pezizomycotina is also immense. Biosynthetic gene clusters (BGCs) encode genes responsible for secondary metabolite production, with the number of BGCs corresponding to the number of secondary metabolites an organism produces. The ∼85,000 known species of Pezizomycotina are estimated to contain 2.5 to 4.2 million biosynthetic gene clusters (BGCs), which are expected to be responsible for the biosynthesis of a similar number of secondary metabolites^23^. There are numerous Pezizomycotina-derived secondary metabolites in pharmaceutical use, including the antibiotic penicillin and the cholesterol-lowering drug lovastatin, with many more in development^24^. Secondary metabolites are also directly linked to fungal pathogenicity^25,22,26,27^. To date, only ∼30,000 fungal secondary metabolites have been identified, revealing the enormous, untapped chemical diversity that remains to be explored. Further, while compartmentalization of secondary metabolite production in organelles has been shown for a handful of metabolites^28–32^, the cellular location of each enzymatic step as well as how intermediates move from one location to another for most is uncharacterized, highlighting major unexplored areas of eukaryotic cell biology.

The Pezizomycotina fungi also have complex life cycles that are governed by environmental cues^33–35^. For example, in *Aspergillus nidulans* light controls physiological and morphological responses. Light is sensed through the phytochrome FphA^36^, the white collar homologs LreA and LreB, and the photolyase-like cryptochrome CryA^37^. Several studies have shown that red light acts as a repressive signal, shifting development away from the sexual cycle and towards the asexual cycle^37–40^ (Figure 1B). While some of the phytochromes, signaling cascades, and transcription factors involved in light sensing are known, much remains to be learned.

Here, we set out to explore the physiological role of organelle dynamics in *Aspergillus* species and discovered that endosome motility and the presence of PxdA on those endosomes is required for light-responsive fungal development. We also discovered that genes involved in secondary metabolism in *A. nidulans* were significantly upregulated in strains that lacked endosome motility and peroxisome hitchhiking. In both *A. nidulans* and the pathogenic species *A. fumigatus* we found that secondary metabolite production was dramatically altered in the absence of *pxdA*. Our results have important implications for optimizing the production of secondary metabolites in heterologous systems and for understanding mechanisms of virulence, especially given the importance of secondary metabolites in *A. fumigatus* pathogenicity.

## Results

### Endosomal motility is required for light-responsive fungal development

Because PxdA is found specifically within the Pezizomycotina subphylum of fungi^17^, we sought to characterize the role of organelle dynamics in physiology that is unique or expanded in this lineage. The velvet protein VeA is a master regulator of several Pezizomycotina-specific processes including their unique developmental programming and secondary metabolism in response to light^41^. Wild-type “velvet” strains of *A. nidulans* produce distinct reproductive structures depending on environmental conditions; for example, they favor asexual structures (conidiophores and conidiospores) under light conditions and sexual structures (cleistothecia and ascospores) under dark conditions (Figure 1B) ^42^. However, our previous studies on PxdA used attenuated *A. nidulans* strains with a mutated version of the velvet gene known as *veA1,* which typically fail to initiate the sexual cycle, even in darkness, making them preferable for routine handling in the laboratory^43^. We therefore generated organelle motility mutants in a non-attenuated isolate of *A. nidulans* that can respond to light (wild-type “velvet” strain or *veA*+). After generating *hookA*, *pxdA*, and *acbdA* deletion mutants in the *veA*+ background, we quantified peroxisome motility, confirming that peroxisomes do not move in *hookAΔ, pxdAΔ* and *acbdAΔ* strains^44,45^ (Figure 1C and 1D).

We observed a growth defect in *hookAΔ* and *pxdAΔ* strains, but not in *acbdAΔ* strains in the light-sensing *veA*+ background (Figure 1E, 1F, S1A and S1B). This was unexpected because the growth phenotype in non-light sensing *veA1* strains was modest as compared to *veA*+ strain^11^ (Figure S1C-S1F). The wild-type *veA*+ strain produced significantly more asexual spores (conidia) in the light versus the dark (Figure 1G). Strikingly, both *hookAΔ* and *pxdAΔ* strains produced significantly fewer conidia in the light as well as in the dark compared to a wild-type strain (Figure 1G). The *hookAΔ* and *pxdAΔ* strains also produced significantly more sexual fruiting bodies (cleistothecia) in the light compared to wild-type strains, a phenotype that is unexpected since cleistothecia are usually formed in the dark (Figure 1H). Deletion of *acbdA* was not significantly different from the wild type (Figure 1E-H).

Complementation of the *pxdAΔ* strain with a wild-type copy of *pxdA* restored light-dependent repression of sexual development to wild-type levels, confirming that the phenotype observed in the *pxdAΔ* strain is specific to the loss of *pxdA* (Figure 1I-1L). In addition to light, nutrient availability is a major environmental factor influencing developmental fate in *A. nidulans*. For example, glucose starvation represses sexual development and cleistothecia formation^36,37^. To determine if the developmental phenotype we observed for *hookAΔ* and *pxdAΔ* strains was light-, but not nutrient- dependent, we grew our mutant strains under starvation conditions (no glucose) and monitored cleistothecia production. We observed no differences in cleistothecia number across our strains, indicating that the developmental phenotypes we observed are light-dependent (Figure S1G-S1I).

Together, our results suggest that early endosome movement and the early endosome-associated protein PxdA are required for light-responsive reproductive development in *A. nidulans*. In contrast, the peroxisome-associated protein AcbdA is not required for reproductive development, showing that the developmental defect seen in *hookAΔ* and *pxdAΔ* strains is independent of peroxisome hitchhiking.

### Genes involved in development are differentially expressed in the absence of PxdA-marked endosome motility

To further investigate Pezizomycotina-specific physiology and understand how *A. nidulans* adapts to the loss of endosome motility and peroxisome hitchhiking, we performed RNA-sequencing analysis of *hookA*, *pxdA, and acbdA* deletion strains. We also included a prototrophic wild-type strain with a complemented copy of the auxotrophic selection marker as a negative control (prototrophic control from here)^48^. All strains were grown in the dark for 24 hours followed by 96 hours in the light before RNA extraction and sequencing (Figure 2A). Variation among replicates was low as was variation between the wild-type and prototrophic control strain (Figure 2B). We detected reads for 9,423 transcripts in the wild-type strain and 9413, 9537, 9513, and 9441 transcripts from the prototrophic control, *hookAΔ, pxdAΔ,* and a*cbdAΔ,* respectively (Data S1). The number of transcripts that were significantly differentially expressed (adjusted p-value<0.01) in mutants compared to the wild type were 552 (5.9%), 6638 (69.6%), 6476 (68%), and 3046 (32.3%) in the prototrophic wild-type, *hookAΔ, pxdAΔ,* and a*cbdAΔ* strains, respectively (Figure S2A-S2D).

**Figure 2.**
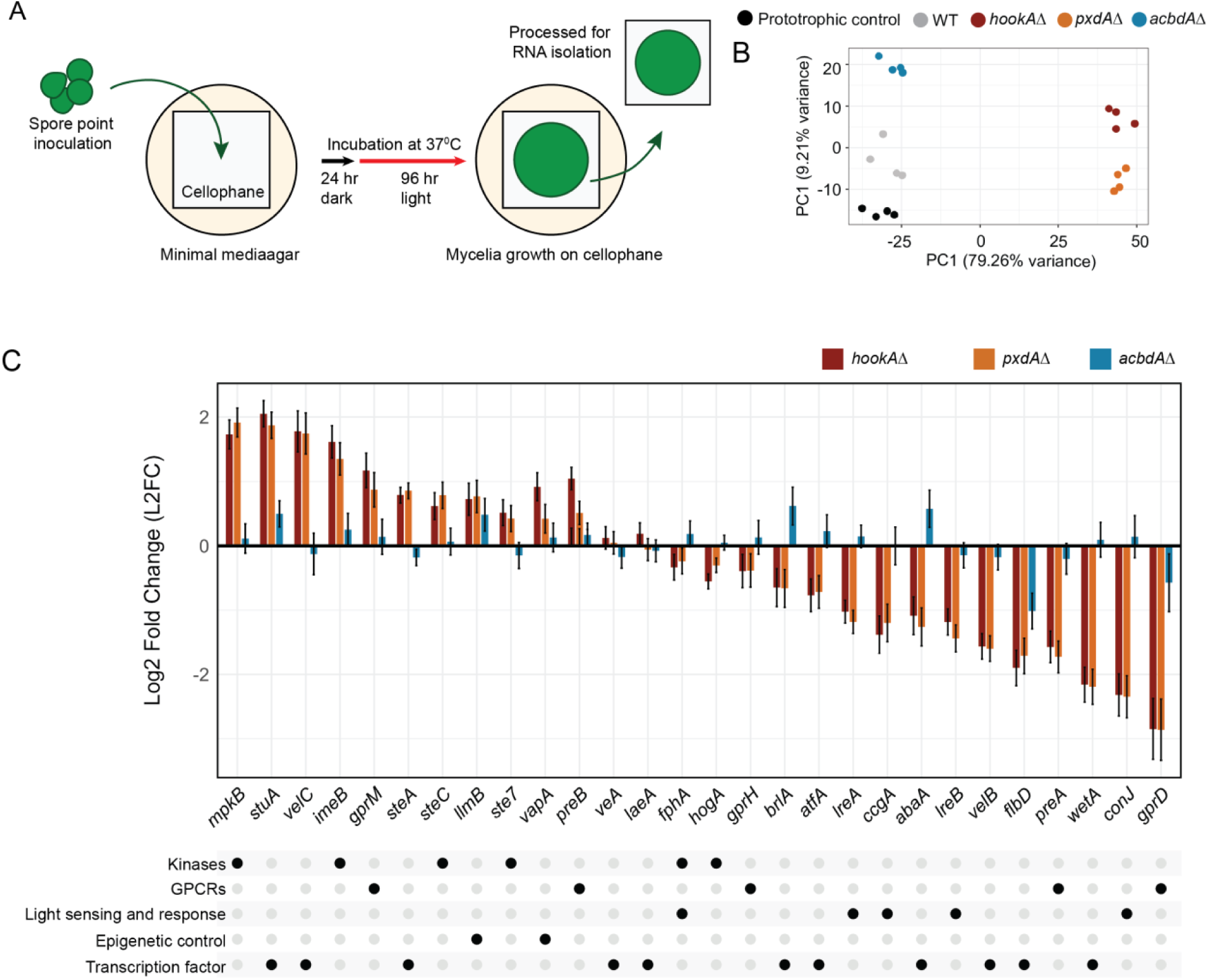
Developmental pathway genes are differentially expressed in the absence of PxdA-marked motile endosomes. **(A)** Schematic depicting the RNA seq experiment design performed with wild-type, prototrophic control, hookAΔ, pxdAΔ, and acbdAΔ strains. (**B**) Principal component analysis of all 20 samples. PC1 explains 79.26% of the variance. Wild-type, the prototrophic control, and acbdAΔ strains group together on the negative end of the axis, while hookAΔ and pxdAΔ strains group together on the positive end. Separation between prototrophic control, wild-type and acbdAΔ strains are observed along PC2, which explains 9.21% of the variance. Some separation between hookAΔ and pxdAΔ strains is also observed along PC2, but to a smaller extent than wild-type and acbdAΔ strains. (**C**) Bar graph of log2 fold-change in gene expression between wild-type and each mutant for a subset of developmental genes. Data represented as mean ± SE (n = 4, biological replicates).

Given the striking developmental phenotype observed in *hookAΔ* and *pxdAΔ* strains, we first examined genes known to be involved in light response and reproductive development (Figure 2C and S3A-C). The development-associated genes with the highest Log_2_ fold expression increase in *pxdAΔ* and *hookAΔ* were positive regulators of sexual development (cleistothecia formation) or repressors of asexual development (*mpkB*, *velC,* and *steC*)^49–51^. In contrast, development associated genes with the greatest Log_2_ fold decrease in expression in *pxdAΔ* and *hookAΔ* were associated with repressing sexual reproduction or promoting of asexual development (*gprD, wetA, lreA*, *lreB*, *flbD* and *velB*)^49–51^ (Figure 2C). Notably, *veA* and *laeA*, which encode transcription factors that govern light-responsive reproductive development^52,53^ were not significantly different in *pxdAΔ* or *hookAΔ*. None of the development-related genes mentioned above were differentially expressed ±1 Log_2_ fold-change in a*cbdAΔ* (Figure 2C).

We next examined additional signaling proteins and transcription factors known to play major roles in developmental decision-making in *A. nidulans* (Figure 2C and Figure S3A-C). Genes associated with activation of sexual development (*gprB, steA, stuA and atfA*)^49,50,54,55^ were upregulated in *pxdAΔ* and *hookAΔ* mutants (Figure S3A). In contrast, genes involved in repression of sexual development, such as *gprD* and *gprH*, as well as genes associated with light response, including *ccgA* and *conJ*, were downregulated^56^ (Figure 2C). Notably, *imeB*, a known repressor of sexual development, was upregulated, while *gprA,* which positively regulates sexual development, was downregulated (Figure 2C), indicating expression patterns that differed from the general trends observed for other developmental regulators^55,57^.

### Genes involved in secondary metabolism are differentially expressed in the absence of endosome motility and peroxisome hitchhiking

We next examined our RNA-sequencing data globally. Most increased and decreased differentially expressed genes were shared between *hookAΔ* and *pxdAΔ* (Figure 3A and Figure 3B), while a smaller group of differentially expressed genes were shared between *hookAΔ*, *pxdAΔ*, and *acbdAΔ* (Figure 3A and 3B). Gene ontology (GO) enrichment analysis of genes with significantly increased expression in all three mutants recovered significant GO terms related to secondary metabolite biosynthesis (Figure 3C), while genes with decreased expression across all three mutants recovered GO terms related to RNA processing (Figure 3D). The top six most enriched GO terms for *pxdAΔ* and *hookAΔ* were also shared and were all related to secondary metabolic processes, including organic heteropentacyclic compounds, toxins, and sterigmatocystin (Figure 3E and 3F).

**Figure 3.**
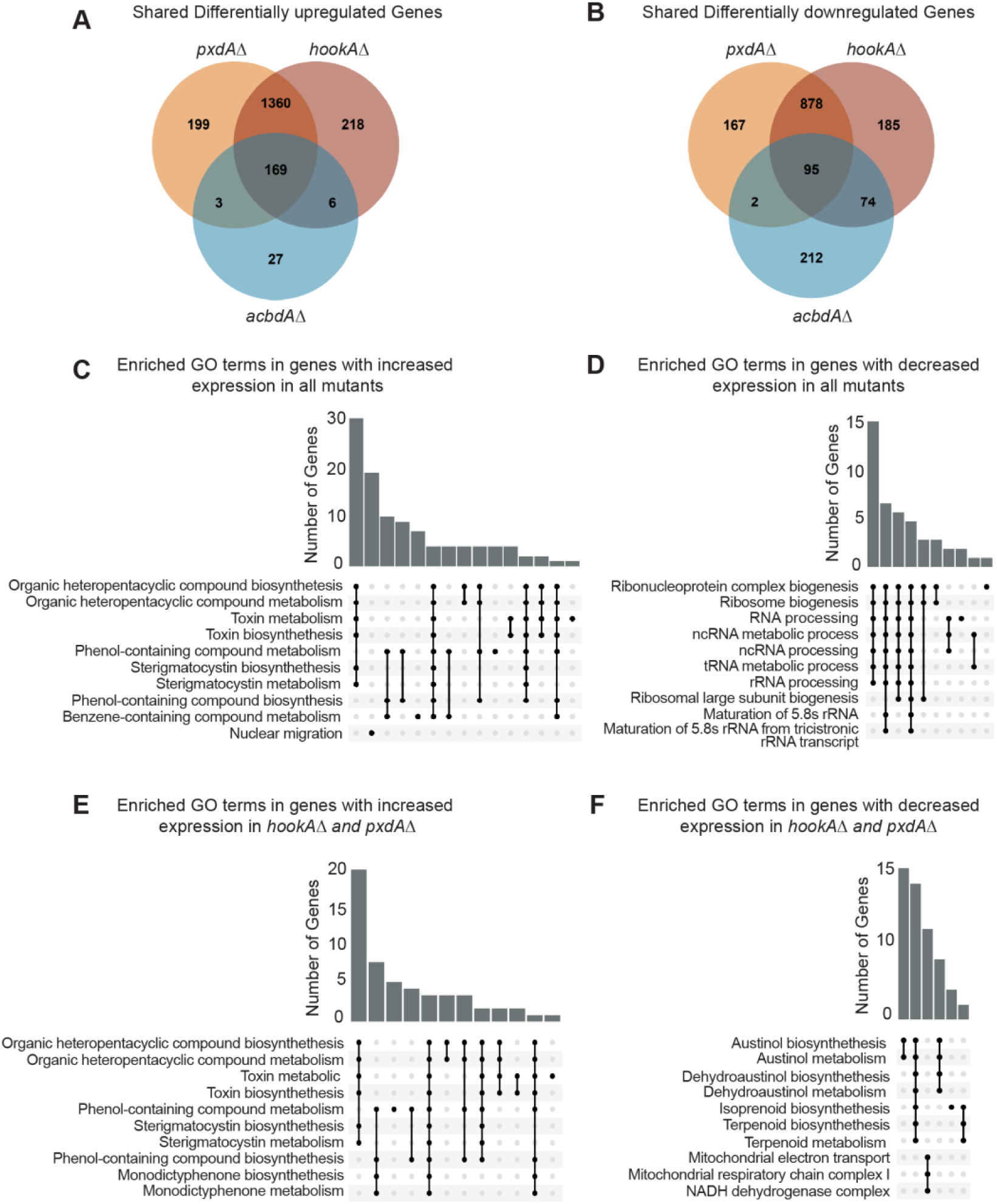
Secondary metabolite genes are differentially expressed in the absence of endosome motility and peroxisome hitchhiking. **(A)** Venn diagram illustrating the number of significantly differentially expressed genes with increased expression in hookAΔ, pxdAΔ, and acbdAΔ strains. (**B**) Venn diagram illustrating the number of significantly differentially expressed genes with decreased expression in hookAΔ, pxdAΔ, and acbdAΔ strains. (**C**) Upset plot of gene ontology enrichment analysis of genes with increased expression in all three mutants. (**D**) Upset plot of gene ontology enrichment analysis of genes with decreased expression in all three mutants. (**E**) Upset plot of gene ontology enrichment analysis of genes with increased expression in both hookAΔ and pxdAΔ strains. (**F**) Upset plot of gene ontology enrichment analysis of genes with decreased expression in both hookAΔ and pxdAΔ strains.

Secondary metabolites are structurally diverse compounds that, unlike primary metabolites, are not directly required for growth. In fungi, the genes for secondary metabolites are organized into biosynthetic gene clusters (BCGs)^58–60^. The biosynthesis of these metabolites involves multiple enzymatic steps and the intracellular transport of intermediates, which requires coordination among organelles such as endosomes, peroxisomes, and mitochondria^31^. Our finding that loss of PxdA-marked endosome motility and peroxisome hitchhiking alters the gene expression of many genes involved in producing distinct secondary metabolites uncovers an unexpected link between organelle dynamics and secondary metabolism.

### PxdA-marked motile endosomes play a role in secondary metabolite production in *A. nidulans*

Given the extensive differential expression of secondary metabolism genes, we next tested whether these changes were reflected in the production of metabolites. *A. nidulans* strains contain ∼60-70 BGCs^61^. Full genome sequencing of our lab strain revealed that it contains a similar number of BGCs (https://doi.org/10.5281/zenodo.18671849). However, only ∼30 secondary metabolites have been characterized in *A. nidulans*^26, 43^. Among the best known are the mycotoxin sterigmatocystin of the aflatoxin family and the antibiotic penicillin. Other identified compounds include austinol, aspernidine A, asperthecin, aspercryptin, nidulanin A, emericellin, emericellamide A–F, emodin, shamixanthone, asperbenzaldehyde, and orsellinic acid derivatives^28,62^.

To directly assess metabolite output, we grew *hookAΔ, pxdAΔ*, *acbdAΔ*, and wild-type strains in light and dark conditions and extracted metabolites for liquid chromatography–mass spectrometry (LC–MS) analysis. The LC–MS profiles revealed substantial alterations in the abundance of several metabolites in *pxdAΔ* and *hookAΔ*, whereas *acbdAΔ* displayed a more modest effect (Figure S4A). Some of these changes aligned with the transcriptional changes from our RNA-seq analysis, whereas others showed the opposite relationship (emericellamide A, emericellamide C/D and asperthecin) (Figure 4A and Figure S4B). To further dissect these effects, we focused on three well-characterized secondary metabolites, sterigmatocystin, austinol, and emericellamide, which also represent distinct biosynthetic classes of fungal secondary metabolites (a polyketide, a meroterpenoid, and a terpene-polyketide hybrid, respectively)^62^.

**Figure 4.**
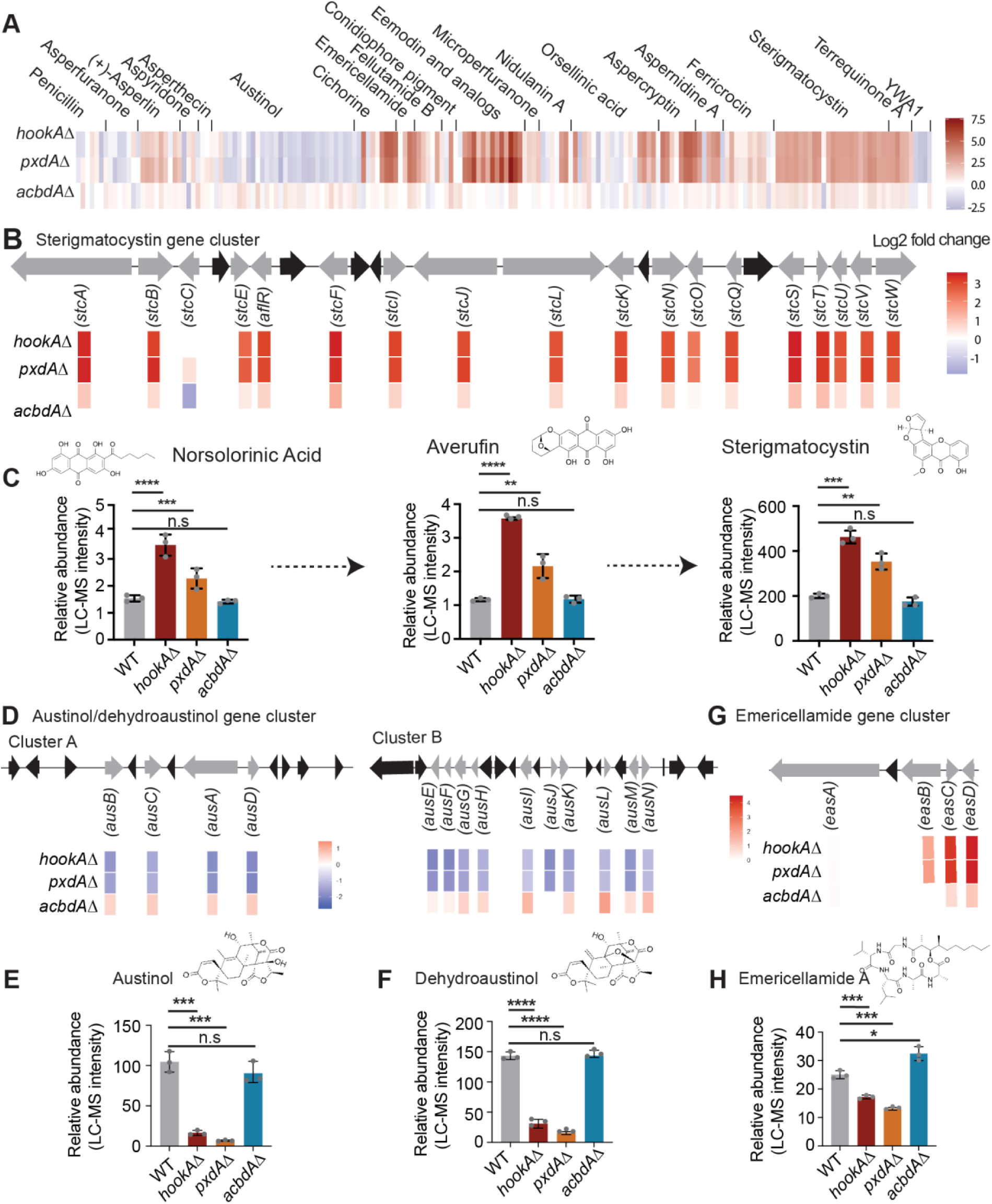
PxdA-marked motile endosomes play a role in secondary metabolite production. **(A)** Heatmap of log2 fold-change in gene expression between hookAΔ, pxdAΔ, and acbdAΔ and wild-type strains for genes involved in secondary metabolism for well-characterized gene clusters that have been associated with a metabolic product. (**B**) Schematic of sterigmatocystin gene cluster (top) and mRNA expression of genes (log2 fold-change vs wild-type) involved in synthesis of sterigmatocystin (bottom). (**C**) Chemical structure of compounds (top) and plots (bottom) of relative abundance of norsolorinic acid, Averufin and sterigmatocystin in wild-type, hookAΔ, pxdAΔ, and acbdAΔ strains (n = 3 biological replicates). (**D**) Schematic of austinol/dehydroaustinol gene cluster and mRNA expression of genes involved in synthesis of austinol/dehydraustinol (top and middle). (E and F) Chemical structures of austinol and dehydrostinol (top) and plots (bottom) of relative abundance of austinol/ dehydroaustinol relative abundance in wild-type, hookAΔ, pxdAΔ, and acbdAΔ strains (n = 3 biological replicates). (**G**) Schematic of emericellamide A gene clusters and mRNA expression (log2 fold-change vs wild-type) of genes involved in synthesis of emericellamide (top). (**H**) Chemical structure of emericellamide A (top) and plot (bottom) of relative abundance in wild-type, hookAΔ, pxdAΔ, and acbdAΔ strains. Data are represented as mean ± SD (n = 3 biological replicates). ** p < 0.01, *** p < 0.001, **** p < 0.0001, n.s (not significant) by unpaired t-test.

Sterigmatocystin is a mutagenic and carcinogenic (Group 2B) mycotoxin that contaminates crop species and exhibits immunotoxic and immunomodulatory activities^63–65^, whose biosynthetic pathway has been extensively studied in *A. nidulans*^66,67^ (Figure 4B). We found increased levels of sterigmatocystin in both the *pxdAΔ* and *hookAΔ* strains, with the *hookAΔ* strain producing significantly higher levels than the *pxdAΔ* strain. In contrast, sterigmatocystin levels in the *acbdAΔ* strain were not significantly different from those in the wild-type strain grown in the light (Figure 4C). The biosynthetic intermediates in the sterigmatocystin pathway, norsolorinic acid and averufin, were also elevated in *pxdAΔ* and *hookAΔ* but unchanged in *acbdAΔ* compared to the wild type (Figure 4C). These results are consistent with our RNA-seq analysis, which showed that nearly all genes within the sterigmatocystin biosynthetic gene cluster were upregulated in the absence of motile endosomes marked by PxdA (Figure 4B and 4C). In contrast, when cultures were grown in the dark for 5 days, sterigmatocystin abundance decreased in *pxdAΔ* but remained comparable to wild type in *hookAΔ* and *acbdAΔ*, despite substantial transcriptional upregulation of the cluster in both *pxdAΔ* and *hookAΔ* mutants in the dark (Figure S4B and Figure S4C).

Austinol and dehydroaustinol belong to the austin-type meroterpenoids, which are known to display antimicrobial and insecticidal activities^68^. In contrast to sterigmatocystin, the levels of both compounds were markedly reduced in the *pxdAΔ* and *hookAΔ* mutants. This decreased production aligned with our RNA-seq analysis, which showed downregulation of all genes in the austinol and dehydraustinol clusters (Figure 4D) in the absence of motile endosomes marked by PxdA. We observed the same trends under both light and dark conditions, although absolute abundances were higher in the light (Figure 4D, 4F, S4D, and S4E).

Emericellamide cyclopeptides, which have been reported to exhibit antibacterial and cytotoxic activities, are produced by several *Aspergillus* species^69^. These compounds showed decreased abundance in *pxdAΔ* and *hookAΔ* mutants under all conditions. However, we observed a slight increase in the abundance of emericellamide in *acbdAΔ* mutants. (Figure 4H and Figure S4F). In contrast, mRNA expression of emericellamide cluster genes was upregulated in all three mutants (Figure 4G and Figure S4B). This decoupling of transcript and product formation has been observed previously^70–72^, suggesting regulation downstream of transcription, potentially involving post translational modifications, metabolite trafficking, precursor availability or subcellular compartmentalization^73,74^. Together, our metabolomic analyses show that disruption of PxdA marked-endosome motility leads to broad changes in secondary metabolism in *A. nidulans*.

### PxdA plays a role in secondary metabolite production in *A. fumigatus*

Having established the role of PxdA in regulating secondary metabolite production in *A. nidulans*, we wondered whether this regulatory role was conserved in the pathogenic fungus *Aspergillus fumigatus*. Although these species diverge in ecology and secondary metabolite repertoires, many core regulatory mechanisms are shared. *A. fumigatus* is an opportunistic pathogen that causes pulmonary and invasive aspergillosis. During infection it produces a variety of secondary metabolites, including helvolic acid, gliotoxin, trypacidin, fumagillin, among others^22^. While *A. fumigatus* strains are predicted to contain ∼40 BGCs^75,76^, only ∼22 secondary metabolites have been characterized^76^.

We began by identifying the *pxdA* ortholog in two well-studied *A. fumigatus* strains (Af293 and CEA10). In silico ortholog prediction identified Afu1g11450 (Af293) and AFUB_010880 (CEA10) as the putative ortholog of *A. nidulans pxdA* (AN1156). The encoded protein, hereafter AfPxdA shares ∼60% protein sequence identity and ∼73% similarity with *A. nidulans* PxdA and shares the same domain structure (Figure 5A). After deleting the gene in both the Af293 and CEA10 backgrounds, we examined the colony morphology of the *AfpxdA*Δ strains. *A. fumigatus* is a heterothallic fungus; it requires partners of two different mating types to initiate sexual reproduction. In the lab a single mating type is grown and thus *A. fumigatus* grows asexually, producing conidia and conidiophores in the presence of both light and dark (Figure 5B). Upon deletion of *AfpxdA*, we observed a reduced colony diameter compared to the wild type in both strains (Figure S5A and S5B). There was no change in the number of conidia made (Figure S5A and S5C) and as expected for a heterothallic fungus, no cleistothecia were produced in either wild-type or *AfpxdAΔ* strains, even in the dark (Figure S5A).

**Figure 5.**
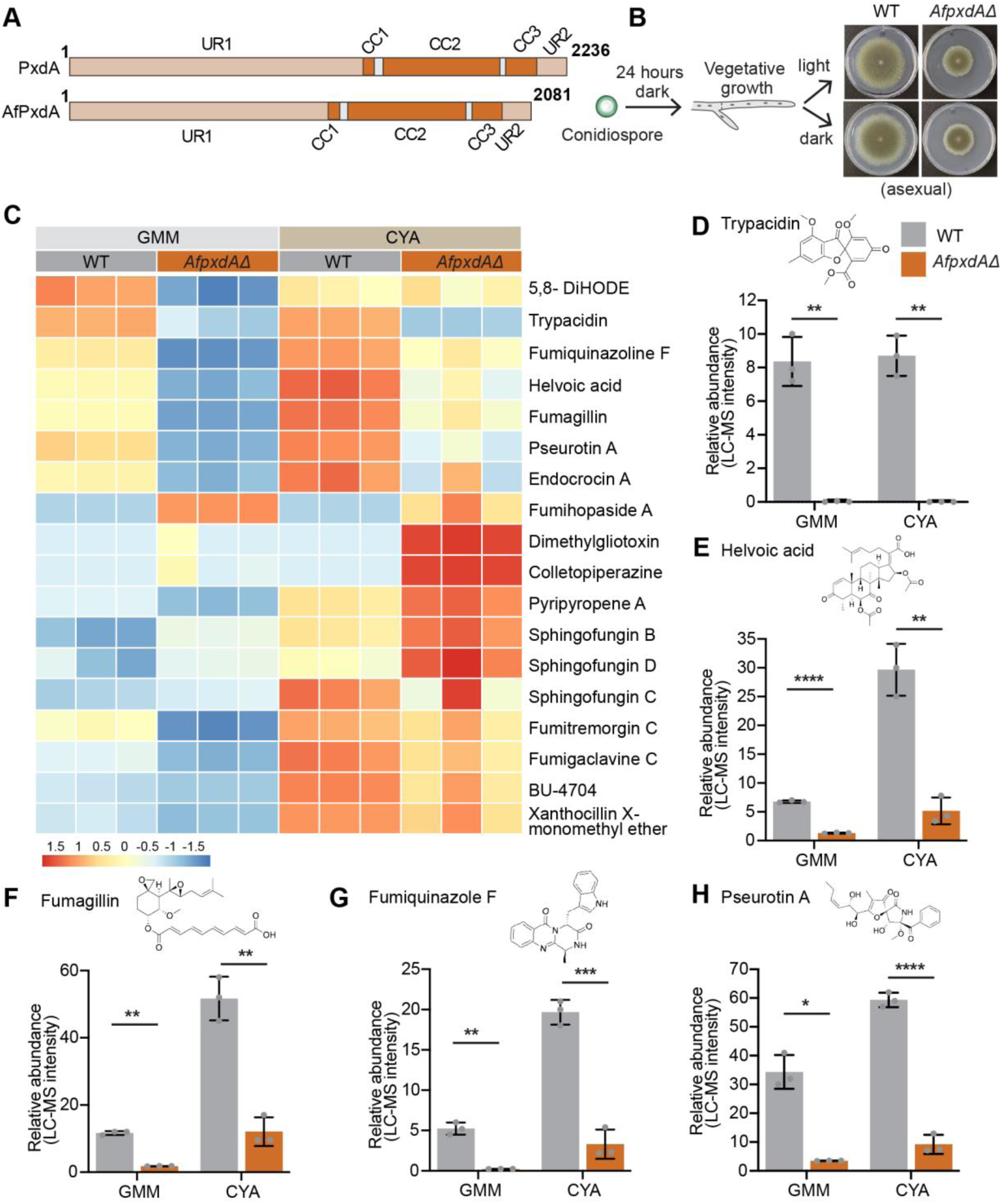
PxdA plays a role in secondary metabolite production in A. fumigatus. **(A)** Schematic of PxdA and AfPxdA domain organization (UR, uncharacterized region; CC, coiled coil). (**B**) Schematic depicting developmental morphology of A. fumigatus wild-type and AfpxdΔ under light versus dark conditions. (**C)** Heatmap showing levels of differentially regulated secondary metabolites in CEA10 wild- type and AfpxdΔ strains grown on GMM and CYA media under dark conditions at 25°C. The color scale represents high (red) and low (blue) abundance values detected by LC–MS analysis. (D-H) Bar plots showing the relative abundance of trypacidin (D), helvoic acid (E), fumagillin (**F**), fumiquinazoline F (**G)** and pseurotin A (H) in wild-type and AfpxdAΔ strains. Data are presented as mean ± SD (n = 3 biological replicates). * p < 0.05, ** p < 0.01, *** p < 0.001 and **** p < 0.0001 by unpaired t test.

Next, we performed LC-MS in wild-type and *AfpxdAΔ* strains in both the Af293 and CEA10 backgrounds. All strains were grown on both glucose minimal medium (GMM) and Czapek yeast extract agar (CYA) (Figure S5A). The LC-MS profiles revealed significant changes in the abundance of several metabolites in both *AfpxdAΔ* strains (Figure 5C and S5D). We identified 18 of the known secondary metabolites in *A. fumigatus*. Of these, five (trypacidin, helvoic acid, fumagillin, fumiquinazoline F, and pseurotin A) were produced at significantly lower levels in both Af293 and CEA10 *AfpxdAΔ* strains (Figure 5D-H and Figure S5D). Dimethyl gliotoxin and colletopiperazine were detected only in Af293, with *AfpxdAΔ* producing significantly less of both metabolites compared to the wild type (Figure S5D). BU-4704, a xanthocillin-like isocyanide was detected in CEA10 grown in CYA media, and its production was significantly less in the *AfpxdAΔ* strain compared to the wild type (Figure 5C). On the other hand, sphingofungin B and sphingofungin D were upregulated in the CEA10 *AfpxdAΔ* strain in both GMM and CYA media, whereas endocrocin was downregulated under both conditions (Figure 5C). Fumihopaside A, dimethylgliotoxin, and colletopiperazine were detected exclusively in the CEA10 *AfpxdAΔ* strain and not in the CEA10 wild-type strain in either medium (Figure 5C). Together, these results indicate that loss of *AfpxdA* profoundly alters the secondary metabolite profile in *Aspergillus fumigatus*.

## Discussion

Organelle positioning and long-range motility are essential for many cellular functions and for maintaining homeostasis in diverse cell types, including filamentous fungi. Here, we investigated the physiological role of organelle dynamics in *Aspergillus* species and discovered that endosome motility and the presence of PxdA on endosomes are required for light-responsive fungal development. We also uncovered an unexpected link between organelle dynamics and the regulation of secondary metabolite gene expression and the production of secondary metabolites. *A. nidulans* strains lacking endosome motility and peroxisome hitchhiking showed upregulation of genes involved in secondary metabolite production. Moreover, in both *A. nidulans* and *A. fumigatus*, loss of the endosomal protein PxdA was associated with altered secondary metabolite profiles. Our findings have important implications for optimizing secondary metabolite production in heterologous systems and for understanding mechanisms of virulence, especially given the importance of secondary metabolites in *A. fumigatus* pathogenicity. Together, our results highlight that organelle dynamics can regulate both fungal development and secondary metabolism.

We discovered that PxdA-marked endosome motility is required for light-responsive reproductive development. Light is a major environmental cue that controls key physiological and morphological responses in fungi^77^. In *A. nidulans,* red-light influences developmental decisions, repressing sexual development. Our finding that *hookAΔ* and *pxdAΔ* strains undergo sexual development even in the presence of red light indicates a defect in light-mediated repression of sexual development. This phenotype mirrors that of *fphAΔ* strains^78^, which lack the red-light phytochrome, raising the possibility that motility of PxdA-marked early endosomes contributes to light-sensing signaling pathways. Roughly 10% of the *A. nidulans* genome is differentially regulated by red light, and most of these genes fail to be induced in *fphAΔ* strains^79^. In our dataset, approximately 60% of red-light-responsive genes were not induced in *hookAΔ* and *pxdAΔ* strains. The overlap in transcriptional changes, together with the shared developmental phenotype between *fphAΔ, pxdAΔ*, and *hookAΔ* strains, suggests that endosome motility plays a role in light sensing in *A. nidulans*. We hypothesize that endosomes transport a signaling factor that is critical for light sensing. While we have not yet identified the specific signaling components transported on motile endosomes, multiple lines of evidence support the role of endosomes as signaling hubs in filamentous fungi. For example, kinases and transcription factors can travel from hyphal tips to nuclei in response to environmental signals^80^, and vesicle-associated signaling has been reported in both *A. nidulans*^81,82^ and *Ustilago maydis*^83^. In *A. nidulans*, the PxdA interactor DipA (a phosphatase) localizes to endosomes and regulates the phosphorylation and degradation of DenA/Den1, a key regulator of asexual development^84^. Finally, PxdA’s conserved N-terminal disordered region contains PAM2L motifs resembling those that mediate RNA hitchhiking in *U. maydis*^10^. Future experiments will identify which proteins and mRNAs require PxdA-marked endosomes to regulate light-responsive development in *A. nidulans*.

Our work also suggests that organelle dynamics contribute to secondary metabolite production in two distantly related *Aspergillus* species. Filamentous fungi produce a diverse array of secondary metabolites that support survival in changing environments. GO term analysis of our mutants defective for endosome and peroxisome motility revealed enrichment for genes involved in secondary metabolite biosynthesis and metabolism, indicating that loss of organelle dynamics broadly influences secondary metabolite gene expression. In *A. nidulans*, *hookAΔ* and *pxdAΔ* mutants showed altered production of sterigmatocystin, austinol, emericellamide, among other metabolites. While we did not observe significant changes in metabolite production in the absence of peroxisome hitchhiking (*acbdAΔ*), the significant transcriptional changes we observed suggest that peroxisome motility is involved in positive and negative feedback loops for secondary metabolite gene expression. In the pathogenic species *A. fumigatus*, deletion of PxdA also led to significant changes in metabolites such as fumagillin and trypacidin, which are implicated in fungal survival and virulence^22^. As the biosynthesis of some secondary metabolites is medium-dependent, it is likely that endosome motility or peroxisome hitchhiking also contributes to the biosynthesis or secretion of metabolites not detected here. Consistent with a broader role for endosome-based regulation of metabolism, endosome motility has been linked to effector production and secretion in *U. maydis*^83,85^ and amylase secretion in *Aspergillus oryzae*^86^. Future experiments will be directed at determining if PxdA-marked endosomes play a signaling role for secondary metabolite production and/ or if endosome motility and organelle hitchhiking is important for metabolite production more directly. For example, some proteins involved in secondary metabolite production are localized to distinct organelle populations^29,28,31,87^, but the role of organelle localization in secondary metabolite production is largely underexplored.

Together, our findings support a model in which HookA-dependent endosome motility and PxdA-marked endosomes contribute to the organization of molecular cues underlying light-dependent developmental regulation and secondary metabolism in filamentous fungi. Although filamentous fungi produce a vast diversity of secondary metabolites, the cellular mechanisms that coordinate their biosynthesis remain poorly understood. Our results suggest that, beyond gene cluster activation, secondary metabolite production depends on spatial and temporal coordination among organelles. Each organelle, including endosomes and peroxisomes, provides distinct biochemical environments that influence enzyme activity, substrate availability, and metabolite stability. How interactions among these compartments are dynamically regulated in response to environmental cues remains an open question. Elucidating these mechanisms will advance our understanding of how organelle dynamics link environmental cues to fungal development and secondary metabolism. These insights are also relevant for interpreting pathogenic mechanisms in filamentous fungi and for guiding strategies to engineer and optimize the heterologous production of medically and industrially important secondary metabolites.

## Material and Methods

### Fungal growth conditions

All strains of *A.nidulans* and *A.fuigatus* were maintained and propagated on solid agar media containing yeast-glucose (YG) complete medium or minimal medium (MM) (Szewczyk et al., 2006). YG and MM agar plates were supplemented with 10 g/L agar-agar or gum agar (USB 10654). YG complete media contained Bacto yeast extract (5 g/L), D-glucose at a final concentration of 2% (20 g/l), trace elements solution (1 ml/L), uracil (56 mg/L), uridine (122.1 mg/L), and riboflavin (2.5 mg/L). The trace elements stock solution (1000×) included: ZnSO₄·7H₂O (22 g/lLH₃BO₃ (5 g/L), MnCl₂·4H₂O (5 g/L), FeSO₄·7H₂O (5 g/L), CoCl₂·5H₂O (1.6 g/L), CuSO₄·5H₂O (1.6 g/L), (NH₄)₆Mo₇O₂₄·4H₂O (11 g/L), and Na₄EDTA (50 g/L). Standard MM was prepared with 1% final D-glucose (10 g/L), MgSO₄ (2 ml/l of 26% wt/vol), trace elements (1 ml/L), and stock salt solution (50 ml/L). The stock salt solution (20×, pH 6.0–6.5) contained NaNO₃ (120 g/l), KCl (10.4 g/L), KH₂PO₄ (30.4 g/l), and MgSO₄·7H₂O (10.4 g/l). For the growth of auxotrophic strains, the necessary supplements were added to the MM at specified working concentrations; pyrG89 allele: 0.5 mM uracil and 0.5 mM uridine; pabaA1 allele: 1.46 μM p-aminobenzoic acid; riboB2 allele: 6.6 μM riboflavin; pyroA4 allele: 3.0 μM pyridoxine. For the light exposure experiments, plates were incubated at 37°C in LED light incubator (Percival Scientific, model: CU-30L2) with the red and blue light set to 50% and 20%, respectively.

### Strain construction

All strains, plamids, and primers used in this study are listed in Table S1, S2 and S3 respectively. All DNA constructs were designed for homologous recombination at the endogenous locus using the endogenous promoter, unless otherwise stated. New plasmid constructs were cloned using Gibson isothermal assembly^88,89^. Fragments were amplified by PCR (Platinum SuperFi II DNA Polymerase, Catalog #12369010) from other plasmids or *Aspergillus* genomic DNA and subsequently assembled into the Blue Heron Biotechnology pUC vector. All plasmids were confirmed by whole plasmid sequencing.

New strains were generated using homologous recombination by transforming PCR-amplified, linearized DNA (2.5 µg) into *A. nidulans* protoplasts to replace the coding sequence in strains lacking *ku70* or *ku80 with AfpyrG (Aspergillus fumigatus pyrG), or Afpyro (Aspergillus fumigatus pyro)*^11,13^. Strain genotypes were verified by PCR amplification of isolated genomic DNA^90^ and Sanger sequencing.

The cloning of *pxdA*-mKate2::*AfpyroA* were described previously^11,14^. To create deletion strains (*hookAΔ*::*Afpyro, pxdAΔ*::*Afpyro*, *acbdAΔ*::*Afpyro*) in *A. nidulans*, two 1 kb DNA fragments directly upstream and downstream of gene of interest, were amplified by PCR from *A. nidulans* genomic DNA, and were subsequently fused to 1.8 kb *Afpyro* fragment and assembled in Blue Heron Biotechnology pUC vector. For cloning of *GFP-acuE:: AfPyrG,* codon-optimized mTag-GFP with a GA(x4) linker was followed by AcuE, its native 3’UTR and an *AfpyrG* cassette, and flanked by 1 kb homologous recombination arms and assembled in Blue Heron Biotechnology pUC vector. To generate the *pxdA* complementation strain (pxdAΔ::pxdA⁺) in the *pxdAΔ::AfpyrG* background of *A. nidulans*, the pxdA gene along with its 3′ UTR was amplified from genomic DNA and fused to the 1.8 kb *Afpyro* fragment, as well as ∼1 kb upstream and downstream flanking regions of pxdA. The final construct was assembled into a pUC vector using Gibson assembly.

To create *pxdA* deletion strains in *A. fumigatus (*Af293) and *A. fumigatus* (CEA10), two 1kb DNA fragments directly upstream and downstream of *AfpxdA*, were amplified by PCR from *Af293* genomic DNA, and were subsequently fused to 2 kb *parapyrG (Aspergillus parasiticus pyrG)* fragment from pJW24^91^ using double joint PCR . Fungal transformation to Af293 or CEA10 was carried out using the method previously outlined^92^. Transformants were confirmed for targeted replacement of the native locus through Southern blotting with *EcoR*I digests for genomic DNA of wild type (Af293 or CEA10) and transformants with P-32 labeled 5′ and 3′ flanks of the knockout construct to get *AfpxdA* Knockout. DNA extraction, restriction enzyme digestion, gel electrophoresis, blotting, hybridization, and probe preparation were performed by standard methods.

### Hyphal tip peroxisome positioning

Imaging to quantify peroxisome positioning along the hyphae was performed at room temperature using a Nikon Ti2 microscope equipped with a An Apo TIRF 100×/1.49 NA objective (Nikon), Yokogawa W1 spinning-disk confocal scan head and two Prime95B cameras (Photometrics). All images were acquired in 16-bit format. The microscope was controlled using NIS-Elements Advanced Research software (Nikon), and the 488-nm laser from a six-line LUN-F-XL laser engine (405, 445, 488, 515, 561, and 640 nm) was used for imaging. Stage movement in the x, y, and z directions was controlled by an ASI MS-2000-500 stage controller (Applied Science Instruments). Live adult hyphae grown for 14–16 hours on agar plates were imaged as Z-stacks with a step size of 0.2 μm spanning a total depth of 8–10 μm. Maximum-intensity projections were generated from the fluorescence images, and the corresponding bright-field images were used to trace the hypha from the tip inward using the segmented line tool (pixel width of 20) in ImageJ/FIJI (National Institutes of Health, Bethesda, MD). These traces were overlaid onto the fluorescence projections to obtain the average fluorescence intensity along the hypha. The mean background intensity for each cell was measured and subtracted from the corresponding line-scan values. Line profile plot was generated using GraphPad Prism version 10 for Mac (GraphPad Software, San Diego, CA, USA; www.graphpad.com).

### Quantification of conidial and cleistothecial development

Asexual and sexual developmental studies were conducted using *Aspergillus nidulans* strains RPA1627, RPA1732, RPA1736, and RPA1756. A total of 15,000 spores were spotted onto MM plates and incubated in the dark at 37 °C for 24 hours. For asexual development, after 24 hours, the plates were transferred to a red-light incubator for the next 4 days. For sexual development, the plates were wrapped in silver foil and kept in the dark at 37 °C for 5 days. After this 5-day period, plates were imaged using an Olympus SZX7 dissection microscope equipped with an Olympus DP28 camera. Images were acquired using Olympus CellSense software and subsequently cropped and processed in Adobe Photoshop 2025.

To quantify conidia, an agar block was excised and homogenized in 0.01% Tween-80, then counted using either a hemocytometer or a u-Count device. For cleistothecia counts, a similarly sized agar block was taken and examined under a dissection microscope. To ensure clear visualization of the cleistothecia, the agar block was washed with 100% ethanol prior to counting. Three agar blocks were taken from each sample for quantification in each experiment, and the experiment was performed at least three times. All values are normalized to wild type strain. Statistical analyses were performed using GraphPad Prism version 10 for Mac (GraphPad Software, San Diego, CA, USA; www.graphpad.com). Graphs were created using the same software. Error bars represent ± SD. Statistical significance is indicated as *P< 0.05, **P< 0.01, ***P< 0.001, ***P< 0.001 and n.s (not significant).

### Secondary Metabolite Extraction

Different strains of *Aspergillus nidulans* were point-inoculated with 1.5x10^4^ spores onto 25 ml of GMM agar plates and allowed to grow at 37 °C for 5 days under the dark and red-light conditions. In the same line, different strains of *Aspergillus fumigatus* were cultured on 25 mL of GMM or CYA agar inoculated with 2.5x10^5^ spores and incubated for ten days at 25 °C in the dark. After the incubation period, each fungal culture along with medium controls (media blanks) were scraped off and transferred into a 50 mL centrifuge tube. The centrifuge tubes were then placed at −80 °C overnight. Next, the entire contents were freeze-dried and lyophilized. Each sample was then extracted overnight with 10 mL or 25 mL of methanol (Sigma Aldrich, St. Louis, MO, USA) for *A. nidulans* and *A. fumigatus* culture, respectively, with shaking at 120 rpm. The extracts were vacuum-filtered, and the solvent was evaporated under reduced pressure. Final extracts were stored at −20 °C. Prior to UHPLC–HRMS analysis, each extract was dissolved in HPLC-grade methanol to a concentration of 2 mg/mL, sonicated until fully dissolved, and filtered using a 0.2μm syringe filter.

UHPLC–HRMS analysis was performed using a Thermo Fisher Scientific Vanquish UHPLC system coupled to a Thermo Q-Exactive Plus mass spectrometer (Thermo Fisher Scientific, Waltham, MA, USA), collecting both MS1 and MS/MS spectra, using an Acquity BEH C18 column (2.1 × 100 mm, 1.7 µm). For *A. nidulans*, samples were analyzed in positive ionization modes across an *m*/*z* range of 188–1800, with an injection volume of 5 μL. The gradient program started at 10% organic for 5 minutes, followed by a linear increase to 90% organic over 20 minutes, then to 98% organic over 2 minutes. The gradient was held isocratically at 98% organic for 5 minutes, decreased back to 10% organic over 3 minutes, and held at 10% organic for an additional 2 minutes, resulting in a total run time of 37 minutes. UHPLC–HRMS RAW files were converted to mzXML format in centroid mode using RawConverter (v1.2.0.1; The Scripps Research Institute, San Diego, CA, USA) and visualized by using the MAVEN2 software (Calico Life Sciences, South San Francisco, CA, USA). Metabolite peaks were confirmed by matching their exact masses and retention times with those of known standards. Except for norsolorinic acid, emericellamides and austinolide, the SMs were predicted based on the comparisons of MS/MS fragmentation patterns with the database and reported data^68^. The peak intensity of each SM was quantified, and relative abundance was calculated by normalizing to an intensity of 1×10⁶, set as 1.

For UHPLC–HRMS with *Aspergillus fumigatus,* samples were analyzed using the equipment previously mentioned under the following method: The chromatographic method consisted of a 1-min isocratic step at 10% MeCN in H2O (both solvents containing 0.05% formic acid), followed by a linear gradient from 10% to 100% MeCN over 20 min, and a 2.5-min wash at 100% MeCN. The flow rate was 0.3 mL/min throughout the run. Data were acquired in both positive and negative ion modes over an m/z range of 100–1500, using higher-energy collisional dissociation (HCD) with a Top 5 data dependent MS/MS method. Raw LC–MS/MS data were converted to .mzML format using MSConvert with peak-picking applied to both MS1 and MS2 levels^94^. Data processing was conducted in MZmine (v4.7.8)^95^. Mass detection used the lowest signal factor, with noise thresholds of 5.0 for MS1 and 2.5 for MS/MS. Chromatograms were built independently for positive and negative modes using a minimum of 4 consecutive scans, a minimum consecutive scan intensity of 1x10^5^, a minimum peak height of 5x10^5^, and a 10 ppm mass tolerance. Detected chromatographic features were processed using Savitzky–Golay smoothing (window: 5 scans), the Local Minimum Feature Resolver, 13C isotope filtering, Join Aligner, Feature Finder, and duplicate peak filter. The final feature list contained 910 features (positive mode) and 1358 features (negative mode). Known Aspergillus fumigatus metabolites were queried based on their expected m/z values ([M+H]+ or [M–H]−) with a 5 ppm tolerance.

Metabolite abundances were represented as the area under the curve of the corresponding chromatographic features. The identities of fumiquinazoline F, fumitremorgin C, and fumagillin were confirmed by comparison with authentic standards. Helvolic acid, pseurotin A, and fumigaclavine C were annotated by matching to gold reference spectra in the GNPS library (https://gnps.ucsd.edu/ProteoSAFe/gnpslibraryspectrum.jsp?SpectrumID=CCMSLIB00011906570#%7B%7D, https://gnps.ucsd.edu/ProteoSAFe/gnpslibraryspectrum.jsp?SpectrumID=CCMSLIB00011906568#%7B%7D, https://gnps.ucsd.edu/ProteoSAFe/gnpslibraryspectrum.jsp?SpectrumID=CCMSLIB00012446712#%7B%7D). Sphingofungin B and endocrocin were annotated by manual comparison with previously published MS/MS data^96^. BU-4704 and fumihopaside A were putative identified based on their predicted molecular formulas and manual validation of their fragmentation patterns (assisted by SIRIUS 5.8.6)^97^. The peak intensity of each SM was quantified, and relative abundance was calculated by normalizing to an intensity of 1×10⁶, set as 1.

### Visualizing secondary metabolite production

For bar plots: statistical analyses were performed using GraphPad Prism version 10 for Mac (GraphPad Software, San Diego, CA, USA; www.graphpad.com). Graphs were created using the same software. Error bars represent ± SD. Statistical significance is indicated as *P< 0.05; **P< 0.01; ***P< 0.001; ns, not significant.

For heatmaps: secondary metabolite quantities were log-transformed then visualized using pheatmap v1.0.13. Metabolites were clustered based on Euclidean distances of their expression profiles across samples, and values were scaled by the pheatmap function to more easily visualize the differences among samples.

### Reference genome generation and analysis

Whole-genome sequencing was conducted for all strains used in this study. Colonies were grown for 16-32 hours in liquid MM+supplements in the dark at 37°C. Tissue was removed from media and extra media was removed via sterile miracloth. Tissue disruption was accomplished by placing tubes with tissue in liquid nitrogen then macerating the fungal tissue with sterile micropestles. Then, 750 μL of lysis buffer (50mM Tris-HCL, 50 mM EDTA, 1% SDS) was added and the tubes were incubated at 60°C for 30 minutes. 375 μL of neutralizing buffer P3 (Qiagen) was added to each tube, which was then mixed and placed on ice for 10 minutes before a 10-minute centrifugation step at room temperature was completed. Supernatant was pipetted onto a filter from the Qiagen Plant Pro DNA extraction kit and was washed according to the manufacturer’s protocol with W1 and W2. The resulting DNA was eluted into 100 μL of water. To clean and concentrate the resulting extraction Mag-Bind TotalPure NGS magnetic beads (omega BIO-TEK, Norcorss, Georgia, USA) were added at 80-100% of the elute volume and cleaning proceeded according to the manufacturer’s protocol. Library preparation was conducted with the Ligation Sequencing gDNA Native Barcoding Kit 24 v14 (SQK-NBD114.24) and resulting libraries were run on R10 flow cells in the Oxford Nanopore Technologies MinION Mk1B (Oxford, United Kingdom). Strains = RPA1732, RPA1736, RPA1756, and 99 were run on another flowcell. Sequences of RPA1627 was generated in single-sample sequencing runs using the LSK-114 library preparation kit.

The multiplexed sequencing run yielded ∼8 Gb of sequence data and sequencing RP 1627 yielded 1.5 Gb of sequence data in binary alignment map format. Base calling and demultiplexing was completed using dorado. Whole *de novo* genome assemblies were created with hifi-asm. BUSCO with Aspergillus odb10 was used to assess the number of contigs, assembly size, contig N50, and the number of complete, single-copy Busco genes. Liftoff was used to annotate each genome with genes from FGSCA4. For each genome the expected assembly insertion was assess using blastn of the entire assembly sequence. The results were used to confirm the length, identify, and completeness of the insert. We also checked the number of copies of GFP present in the genome by using blastx. All code for whole genome sequencing analyses are available at public GitHub repository (https://github.com/jallen73/Aspergillus_genotyping_with_ONT).

### RNA sequencing

Five different strains were used to assess transcriptional changes associated with loss of *hookA, pxdA,* and *acdbA*. We used two different controls, the *veA+* wildtype (RPA1492) and an auxotrophic control differing from the wildtype only in the insertion of a complemented selection cassette (RPA1627). Three mutants with RPA1627 as the control strain were used: *hookAΔ* (RPA1736)*, pxdAΔ* (RPA1732), and *acbdAΔ* (RPA1756). Strains were grown on minimal media with 1% glucose. Sterilized cellophane wrap (Purple Q Crafts) was placed in the center of each petri dish and were point inoculated with equal numbers of spores from each strain. They were grown in the dark at 37 °C for 24 hours then transferred to red light conditions at 37 °C and grown for an additional 96 hours. Saran wrap pieces were removed with sterile forceps and transferred to 1.5 mL Eppendorf tubes. Total RNA was extracted using the RNeasy Mini Kit (Qiagen, Maryland, USA). Initial tissue disruption was accomplished with glass beads and bead beater homogenizer. Total RNA yields were quantified with nanodrop ranged from 1ug-3ug/μl. 150 base pair, paired end sequencing was conducted by Novagene on the Illumina NovaSeqXPlus platform yielding 69 GB of data.

### RNA-Seq transcript quantification

All code for RNA-Seq analyses is deposited on GitHub (https://github.com/jallen73/RNA-seq-Aspergillus-nidulans). Raw fastq quality was checked with fastqc v0.12.1. To trim low quality bases and reads we used FastP v0.24.0 with base correction implemented for overlapping nucleotides, a minimal acceptable phred score of 10 and a read length cutoff of 50 basepairs^98^. Fastqc was used to check that trimming was successful. To quantify the relative abundances of transcript we used salmon v1.10.3^98^. First, FungiDB-65_AnidulansFGSCA4_AnnotatedTranscripts.fasta were indexed. The index was then used with the salmon quant function with the mapping validation, GC bias correction, and sequencing bias options implemented. The abundances of the transcripts for knocked-out genes were verified for each mutant. In RPA1732, *pxdAΔ*, *pxdA* transcripts were recovered at high abundances. To further investigate this finding we aligned all of the reads to the reference genome using bwa v0.7.18^99^, sorted the reads using samtools v1.2^100^, then viewed the aligned transcripts in the Integrated Genome Viewer v2.19.1^101^. The transcripts identified as *pxdA* mapped to the 5′ untranslated region of the gene and did not map to any translated portions of the gene. Thus, we manually set the abundances of these transcripts to zero for the *pxdAΔ* samples before conducting downstream analyses.

### Differential gene expression analyses in R

Analyses were run in R version 4.4.1 implemented in RStudio version 2024.04.2+764. DESeq2 v1.44.0 was used to test for differential gene expression^102^. First, a transcript to gene name annotation file was created using FungiDB-68_AnidulansFGSCA4.gff. The transcript to gene annotations, experiment metadata, and salmon quantifications were merged then used as the input for DESeq2. To visualize the number of genes and the degree of differential expression genome-wide we used volcano plots. To confirm that insertion of a selection cassette alone did not significantly change transcript production, we first verified that the wild type and metabolic control were not significantly different. We examined volcano plots and principal components analysis results to confirm that the metabolic control and the wild type were more similar to each other than to any of the KO strains.

To determine which genes shared significant overexpression and underexpression between and among mutants, gene lists were created by filtering all expression results to include only genes with an adjusted p-value equal to or less than 0.01 and a Log_2_ fold-change greater than one or lower than negative one. Then, venndetail v1.20.0 was used to determine which genes in the list were shared by which mutants and the results were visualized in a venn diagram. The list of genes that were overexpressed or underexpressed in all three mutant strains and in two mutant strains, *pxdAΔ* and *hookAΔ*, were subjected to a gene ontology enrichment analysis using all ontologies, a Benjamini-Hochsberg correction for multiple tests, and a p-value and q-value cutoff of 0.05. Results were visualized with upset plots. The gene ontology enrichment database for *Aspergillus nidulans* was created with the GO.db v3.19.1 and AnnotationForge v1.46.0 libraries and the FGSC A4 reference genome. After removing null values, mapping outdated GO terms to updated terms ‘makeOrgPackage’ was used to create the GO database. To examine the changes in expression level of a suite of development-related genes we calculated mean Log_2_ fold-change and 95% confidence intervals and displayed them in a barplot.

We created heatmaps of additional genes grouped by function: asexual development, balancing sexual and asexual development, germination, light response, sexual development, and vegetative growth. The same function was used to create heatmaps of secondary metabolite biosynthetic gene clusters.

### Figure preparation

Figures were prepared in Adobe Illustrator.

## Resource availability

### Lead contact

Further information and requests for resources and experimental data should be directed to and will be fulfilled by the lead contact, Samara Reck-Peterson.

## Material availability

Any material or reagents used in this paper will be available upon request.

### Data and code availability

The RNAseq and genome sequencing datasets generated and analyzed during this study have been deposited on Zenodo; The DOI is: https://doi.org/10.5281/zenodo.18671849.

The UHPLC–HRMS data generated and analyzed during this study have been deposited on MassIVE (Centre for Computational Mass Spectrometry, UCSD, CA); The DOI is: doi:10.25345/C5PR7N72X. All analysis scripts used for RNA seq (https://github.com/jallen73/RNA-seq-Aspergillus-nidulans) and whole genome sequencing (https://github.com/jallen73/Aspergillus_genotyping_with_ONT) analyses is deposited on Github.

## Acknowledgments

We thank all the members of Reck-Peterson laboratory and Andres Leschziner for critical feedback on experiments. We thank Anu Singh (Reck-Peterson laboratory) for the illustrations of conidiophores and cleistothecia. We would like to thank Wenjun Zhang (UC Berkley) and Haoran Pang (UC Berkley) for helping us in initial LC-MS experiments. We thank Xin Xiang for helpful advice related to strain construction. S.R.P. is supported by the Howard Hughes Medical institute and NIH grant R35GM141825; P.S. by NIH grant F32GM157937; J.S. by NIH grant 5R35GM15085 and N.P.K. by NIN Grant R35GM156119 and NIH grant R01AI150669.

## Author contributions

S.R.P., G.K. and L.D.S.O. developed the project. G.K., L.D.S.O, and J.L.A. designed the experiments. G.K., J.L.A. and L.D.S.O. performed RNAseq experiments. J.L.A. and L.D.S.O. performed RNA seq analysis. M.A.R. and E.A.R. performed the LC-MS experiments. B.D and J.S. provided the expertise and material for *acbdA* mutants. N.P.K. provided expertise and materials for the LC-MS experiments. G.K. and J.L.A. analyzed the results. S.R.P, G.K. and L.D.S.O. created the illustrations. P.H.S. helped with providing critical reagents. S.R.P., G.K., and J.L.A. wrote the manuscript and all authors helped edit it.

## Declaration of generative AI and AI-assisted technologies

ChatGPT 4 was used to draft some R code.

## Supplemental information

Figures S1-S5

Table S1. Strains used in this study

Table S2. Plasmid constructs used in this study Table S3. Primers used in this study

Data S1. Differentially expressed genes for all mutants Data S2. Secondary metabolite analysis for *A. nidulans* Data S3. Secondary metabolite analysis for *A. fumigatus*

---

**Figure S1.**
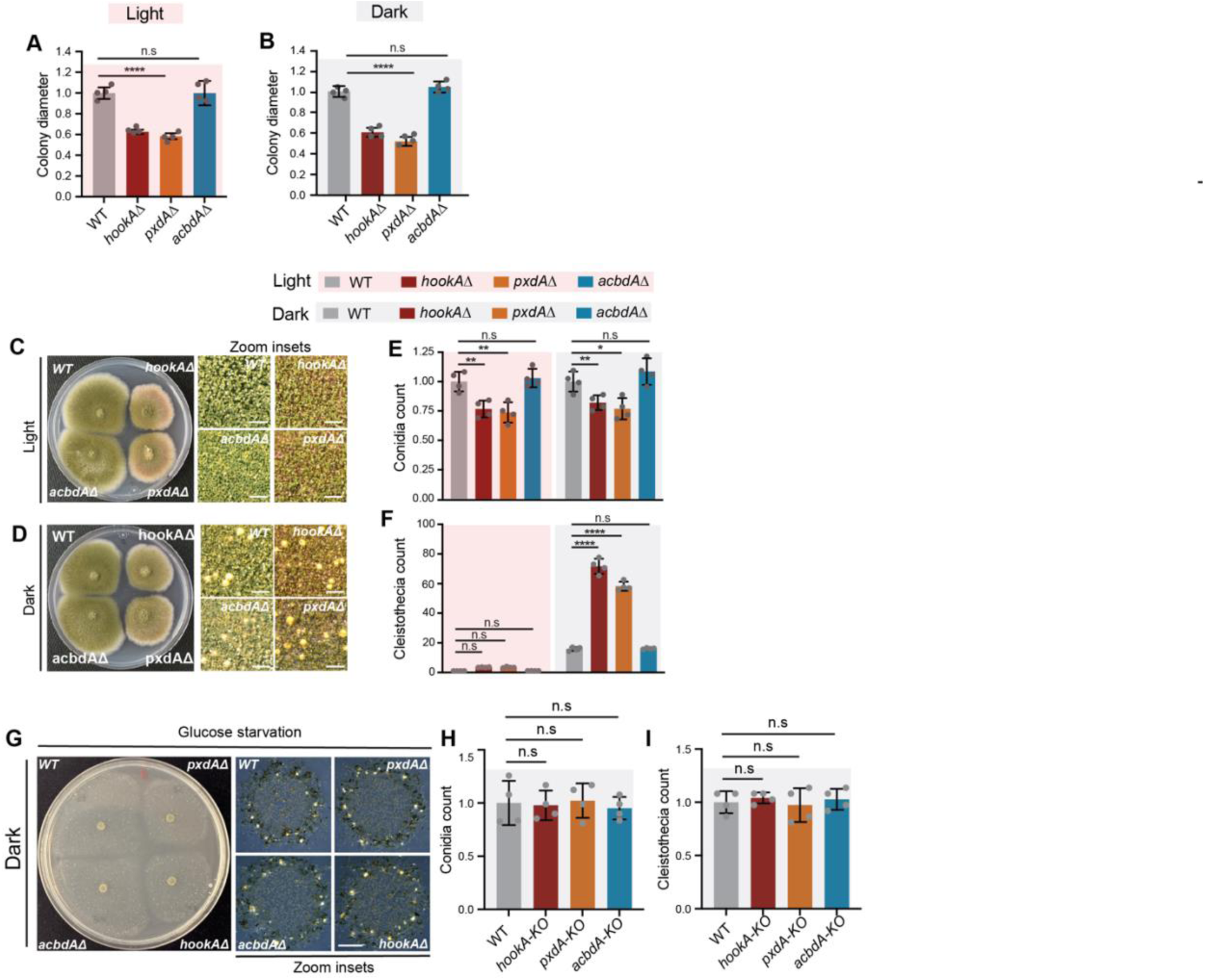
(related to figure 1). Loss of pxdA and hookA reduces radial growth in Aspergillus nidulans. (A and. **B)** Quantification of radial growth in light (**A)** and dark (**B**) condition, normalized to the wild-type strain. Data are represented as mean ± SD (n = 4 biological replicates). **** p < 0.0001, n.s (not significant) by one-way ANOVA. (**C** and **D**) Phenotype for wild-type, hookAΔ, pxdAΔ, and acbdAΔ strains in veA1 background, grown in light (**C)** and dark (**D**). (**E** and **F**) Quantification of conidia (E) and cleistothecia (F) normalized to the wild-type strain grown in light. Data are represented as mean ± SD (n = 4 biological replicates). * p < 0.05, ** p < 0.01, **** p < 0.0001, n.s (not significant) by two-way ANOVA. (**G**) Image of agar plate showing growth phenotype of wild-type, hookAΔ, pxdAΔ, and acbdAΔ strains grown on glucose starvation media under dark condition. (**H** and **I**) Quantification of conidia (**I**) and cleistothecia (**J**) from glucose starvation media plates normalized to the wild-type strain. Data are represented as mean ± SD (n = 4 biological replicates). n.s (not significant) by ordinary one-way ANOVA.

**Figure S2.**
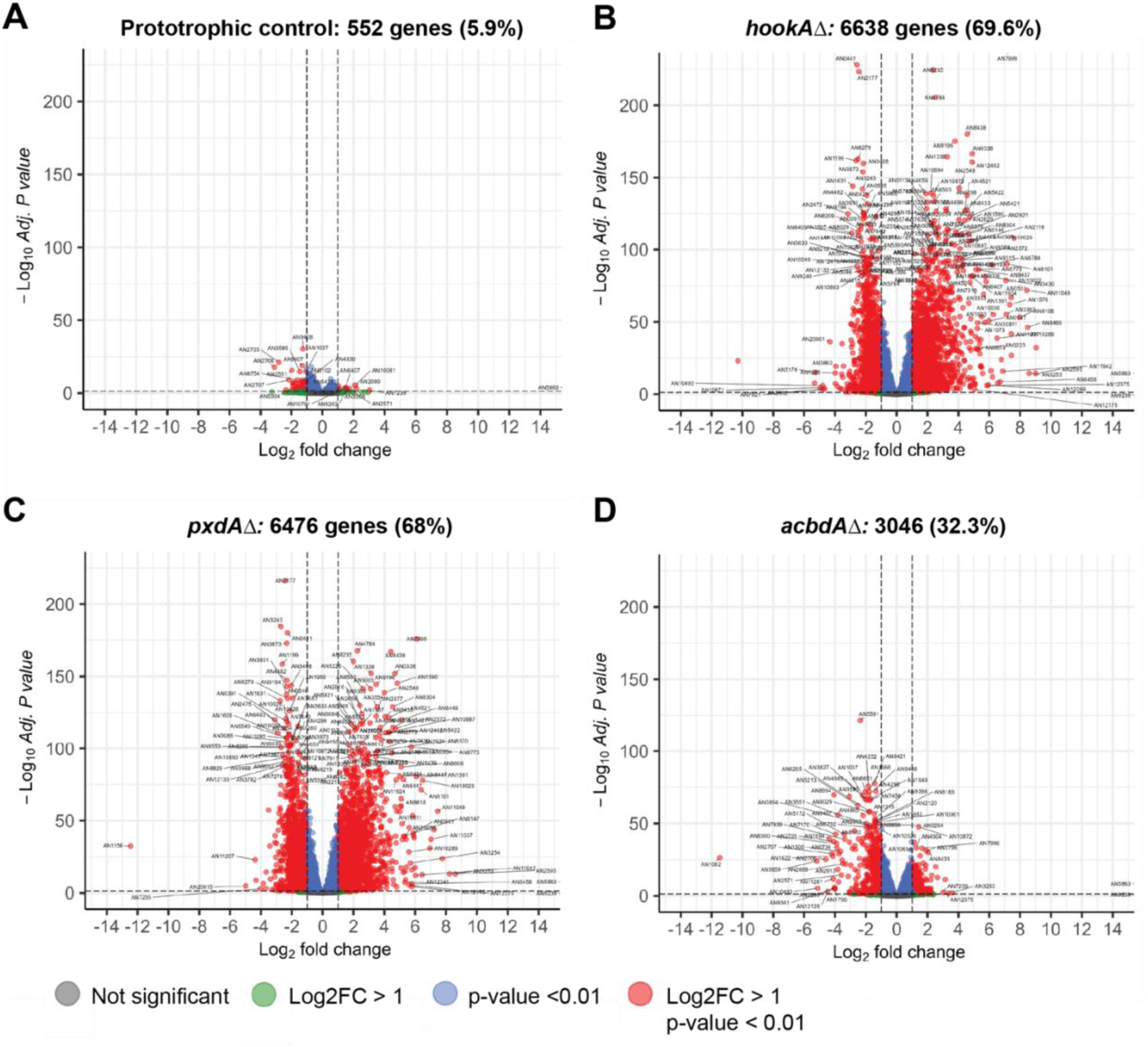
(related to figure 2). Differentially expressed genes in prototrophic control, hookAΔ, pxdAΔ, and acbdAΔ compared to a wild-type strain (A-D) Volcano plots summarizing the Log2 fold-changes in gene expression and adjusted p-values in prototrophic control, hookAΔ, pxdAΔ, and acbdAΔ compared to a wild-type strain. Parenthetical values represent the percentage of detected transcripts that were significantly differentially expressed.

**Figure S3.**
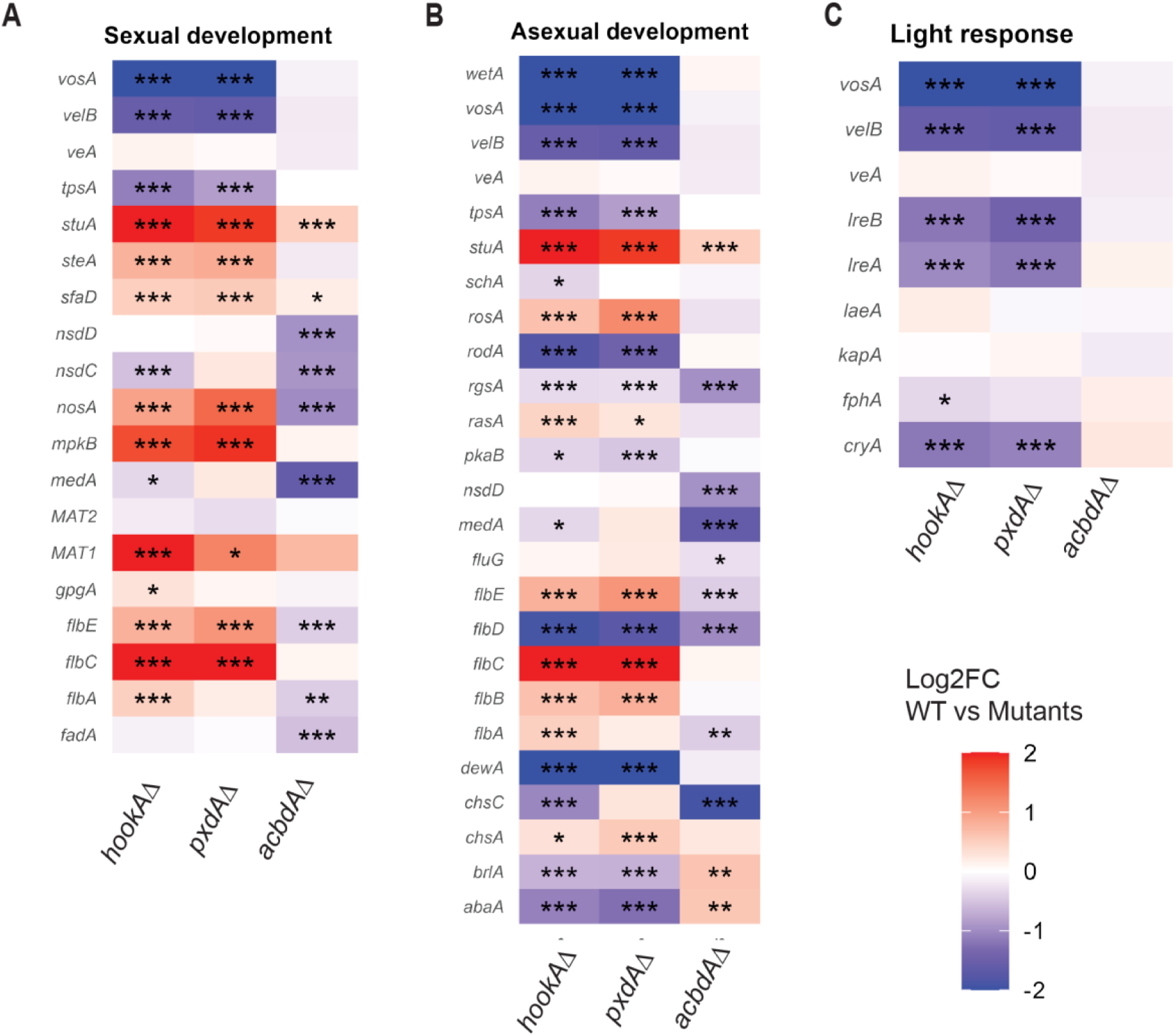
(related to figure 2). Developmental pathway genes are differentially expressed in hookAΔ and pxdAΔ. (A-C) Heatmaps of standardized transcript levels of mutants as compared to the WT for essential reproductive genes involved in asexual development (**A**), sexual development (**B**), and light response (**C**). Data are represented as mean ± SD (n = 4 biological replicates). * p < 0.05, ** p < 0.01, *** p < 0.001, **** p < 0.0001 and n.s. (not significant) by two-way ANOVA.

**Figure S4.**
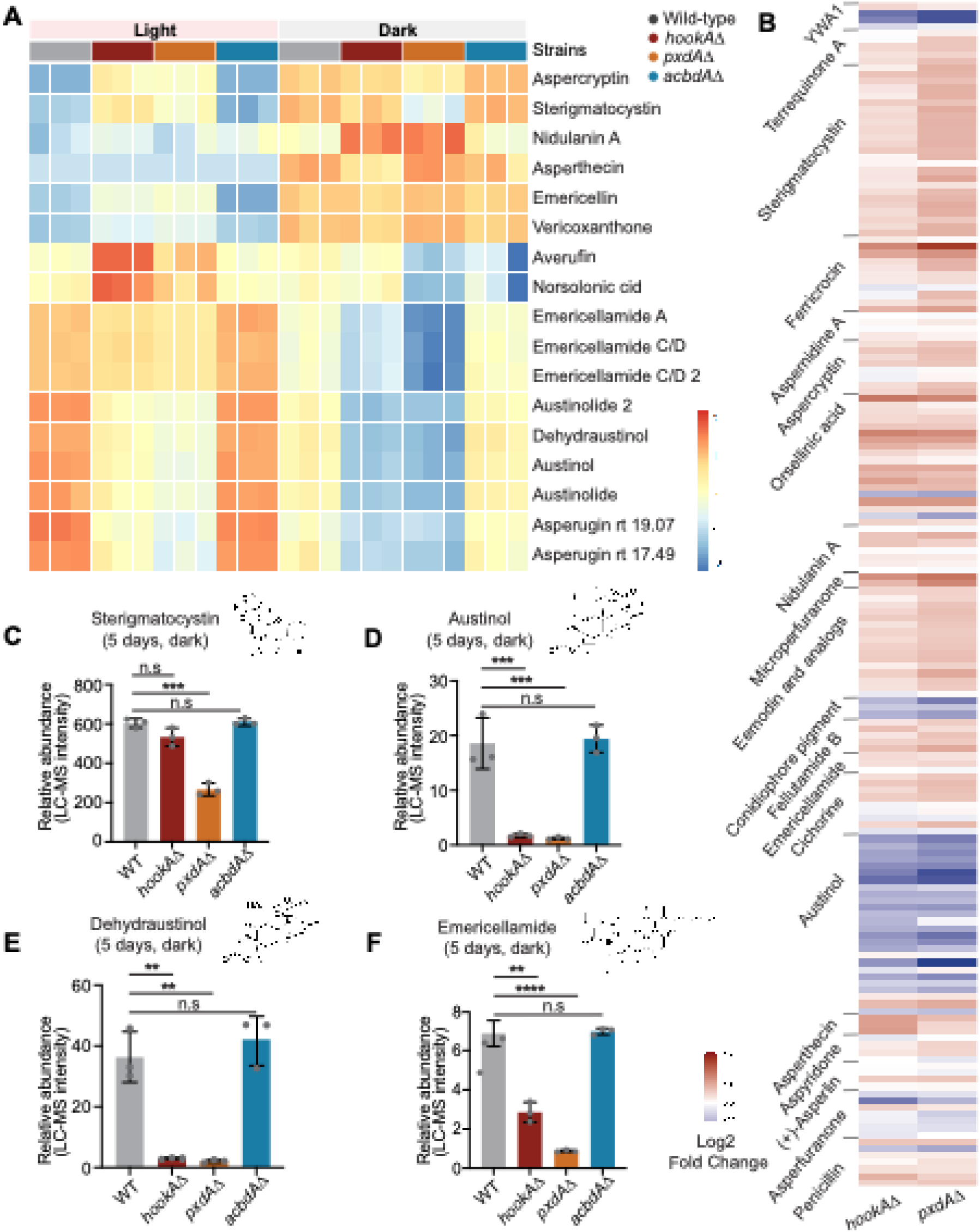
(related to figure 4). Loss of pxdA, hookA, and acbdA alters secondary metabolite gene expression and metabolite production in Aspergillus nidulans. **(A**) Heatmap showing levels of differentially regulated secondary metabolites in WT, pxdAΔ, hookAΔ, and acbdAΔ strains grown under light and dark conditions for five days. The color scale represents high (red) and low (blue) abundance values detected by LC–MS analysis. (B) Heatmap showing log2 fold-changes in gene expression between mutants and wild-type strains grown in the dark for five days, covering all secondary metabolism genes from well-characterized clusters with known products. (**C–F**) Relative abundance plots of sterigmatocystin (**C**), austinol (**D**), dehydroaustinol (**E**), and emericellamide (**F**) in WT, hookAΔ, pxdAΔ, and acbdAΔ strains grown in the dark for five days. Data are represented as mean ± SD (n = 3 biological replicates). ** p < 0.01, *** p < 0.001, **** p < 0.0001 and n.s. (not significant) by unpaired t-test.

**Figure S5.**
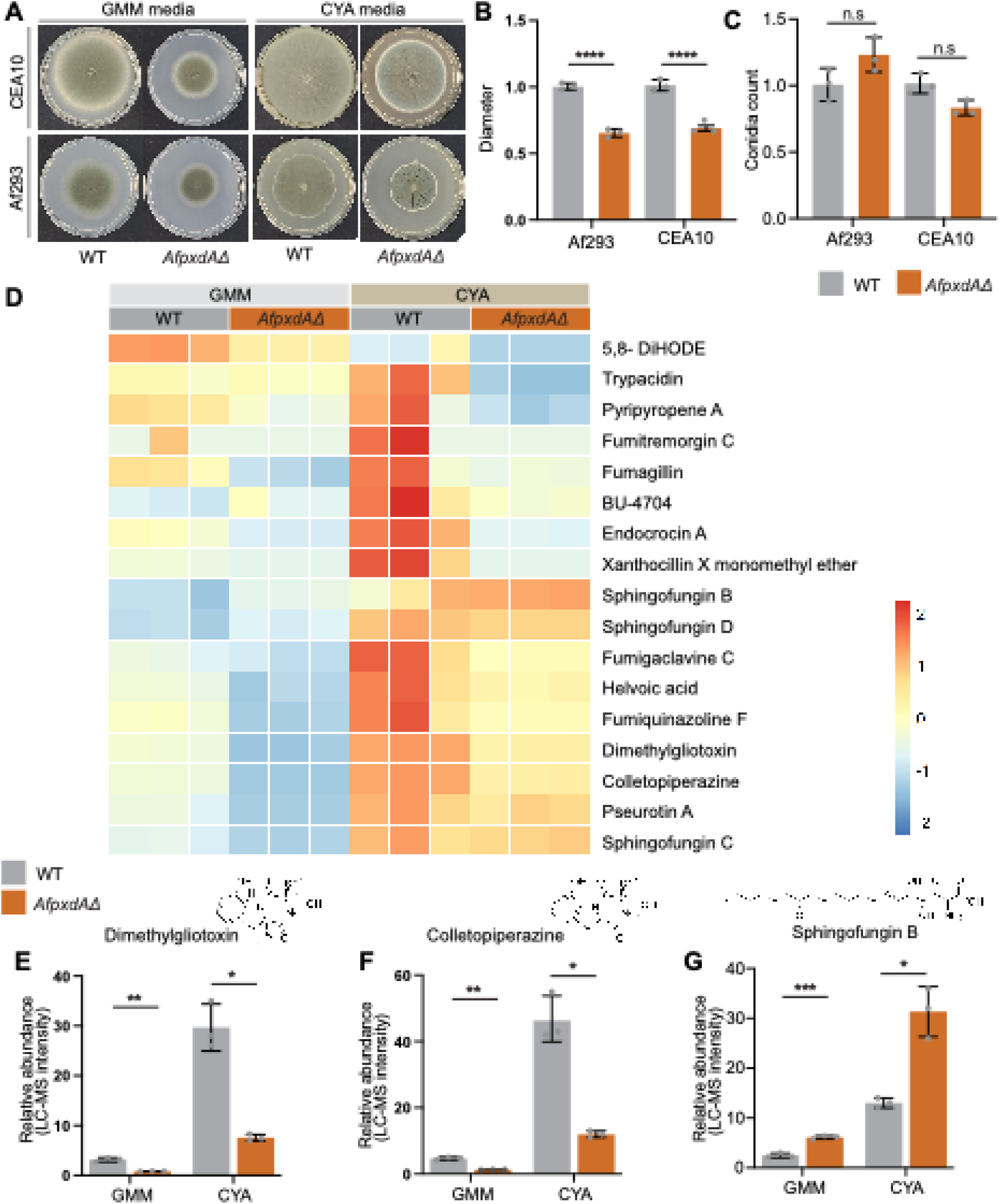
(related to figure 5). Loss of PxdA affects secondary metabolite production in A. fumigatus. **(A)** Image of agar plate showing growth phenotype for A. fumigatus (CEA10 and Af293) wild-type and AfpxdΔ strains grown on glucose minimal media (GMM) and Czapek yeast extract agar (CYA) media under dark conditions. (**B**) Quantification of radial growth on GMM plates normalized to WT strain. Data are represented as mean ± SD (n = 4 biological replicates). ** p < 0.01, *** p < 0.001 and n.s. (not significant) by unpaired t test. (**C**) Quantification of conidia from the strains grown on GMM plates and count was normalized to the WT strain. (D) Heatmap showing levels of differentially regulated secondary metabolites in Af293 wild-type and AfpxdΔ strains grown on GMM and CYA media under dark conditions at 25°C. The color scale represents high (red) and low (blue) abundance values detected by LC–MS analysis. (E-F) Bar plots showing the relative abundance of dimethylgliotoxin (E), colletropiperazine (**F**), in Af293 wild-type and AfpxdΔ strains. (**G**) Bar plots showing the relative abundance of sphingofungin B in CEA10 wild-type and pxdAfΔ strains Data are presented as mean ± SD (n = 3 biological replicates). * p < 0.05, ** p < 0.01 and *** p < 0.001, by unpaired t test.

**Table S1:** Strains used in this study.

| Strain | Genotype | Source |
| --- | --- | --- |
| RPA1492 (SJR2) | *A. nidulans*: *pyrG89, pyroA4, nkuA :: bar, veA<sup>+</sup>* | Gift from Richard Fischer lab |
| RPA1608 | *A. nidulans*: [*mGFP-acuE :: AfpyrG*], *pyrG89, pyroA4, nkuA :: bar, veA<sup>+</sup>* | This study |
| RPA1627 | *A. nidulans*: [*mGFP-acuE :: AfpyrG*], *pyrG89, pxdA-mKate :: Afpyro*, *pyroA4, nkuA :: bar, veA<sup>+</sup>* | This study |
| RPA15 | *A. nidulans*: *pyrG89, pyroA4, nkuA :: bar, veA1* | Deposited in FGSC by Michael Hynes |
| RPA1673 | *A. nidulans*: *pyrG89, [pxdAΔ :: Afpyro]*, *pyroA4, nkuA :: bar, veA1* | This study |
| RPA1675 | *A. nidulans*: *pyrG89, [hookAΔ :: Afpyro]*, *pyroA4, nkuA :: bar, veA1* | This study |
| RPA1732 | *A. nidulans*: [*mGFP-acuE :: AfpyrG*], *pyrG89, [pxdAΔ :: Afpyro]*, *pyroA4, nkuA :: bar, veA<sup>+</sup>* | This study |
| RPA1736 | *A. nidulans*: [*mGFP-acuE :: AfpyrG*], *pyrG89, [hookAΔ :: Afpyro]*, *pyroA4, nkuA :: bar, veA<sup>+</sup>* | This study |
| RPA1756 | *A. nidulans*: [*mGFP-acuE :: AfpyrG*], *pyrG89, [acbdAΔ :: Afpyro]*, *pyroA4, nkuA :: bar, veA<sup>+</sup>* | This study |
| RPA1747 | *A. nidulans*: [*pxdAΔ :: AfpyrG*], *pyrG89, pyroA4, nkuA :: bar, veA<sup>+</sup>* |  |
| RPA1849 | *A. nidulans*: *pyrG89, [acbdAΔ :: Afpyro]*, *pyroA4, nkuA :: bar, veA1* | This study |
| RPA1850 | *A. nidulans*: [*pxdAΔ :: AfpyrG :: pxdA<sup>+</sup> :: Afpyro*], *pyrG89, pyroA4, nkuA :: bar, veA<sup>+</sup>* | This study |
| RPA1832 | *A. fumigatus*: *Af293, ΔnkuA::mluc, pyrG-* | Lim et al. 2018<sup>1</sup> |
| RPA1777 | *A. fumigatus*: *Af293, ΔnkuA::mluc, pyrG-, AfpyrG* | Lim et al. 2018<sup>1</sup> |
| RPA1778 | *A. fumigatus*: *Af293, [pxdAΔ :: parapyrG], ΔnkuA::mluc* | This study |
| RPA1780 | *A. fumigatus*: *CEA10, ku80Δ* | Ferreira ME et al. 2006<sup>2</sup> |
| RPA1781 | *A. fumigatus*: *CEA10, [pxdAΔ :: parapyrG], ku80Δ* | This study |
| RPA1833 | *A. fumigatus*: *CEA10, ku80Δ, pyrg-* | Ferreira ME et al. 2006<sup>2</sup> |

**Table S2:** Plasmids used in this study.

| **Plasmid name** | **Construct name** | **Source** |
| --- | --- | --- |
| RPB2047 | *(mGFP-acuE :: AfpyrG)* | This study |
| RPB990 | *(pxdA-mKate :: Afpyro)* | Salogiannis et al. 2016<sup>3</sup> |
| RPB2646 | *(pxdAΔ :: Afpyro)* | This study |
| RPB2400 | *(hookAΔ :: Afpyro)* | This study |
| RPB2853 | *(acbdAΔ :: Afpyro)* | This study |
| RPB3026 | *(pxdAΔ :: AfpyrG :: pxdA<sup>+</sup> :: Afpyro)* | This study |

**Table S3:**
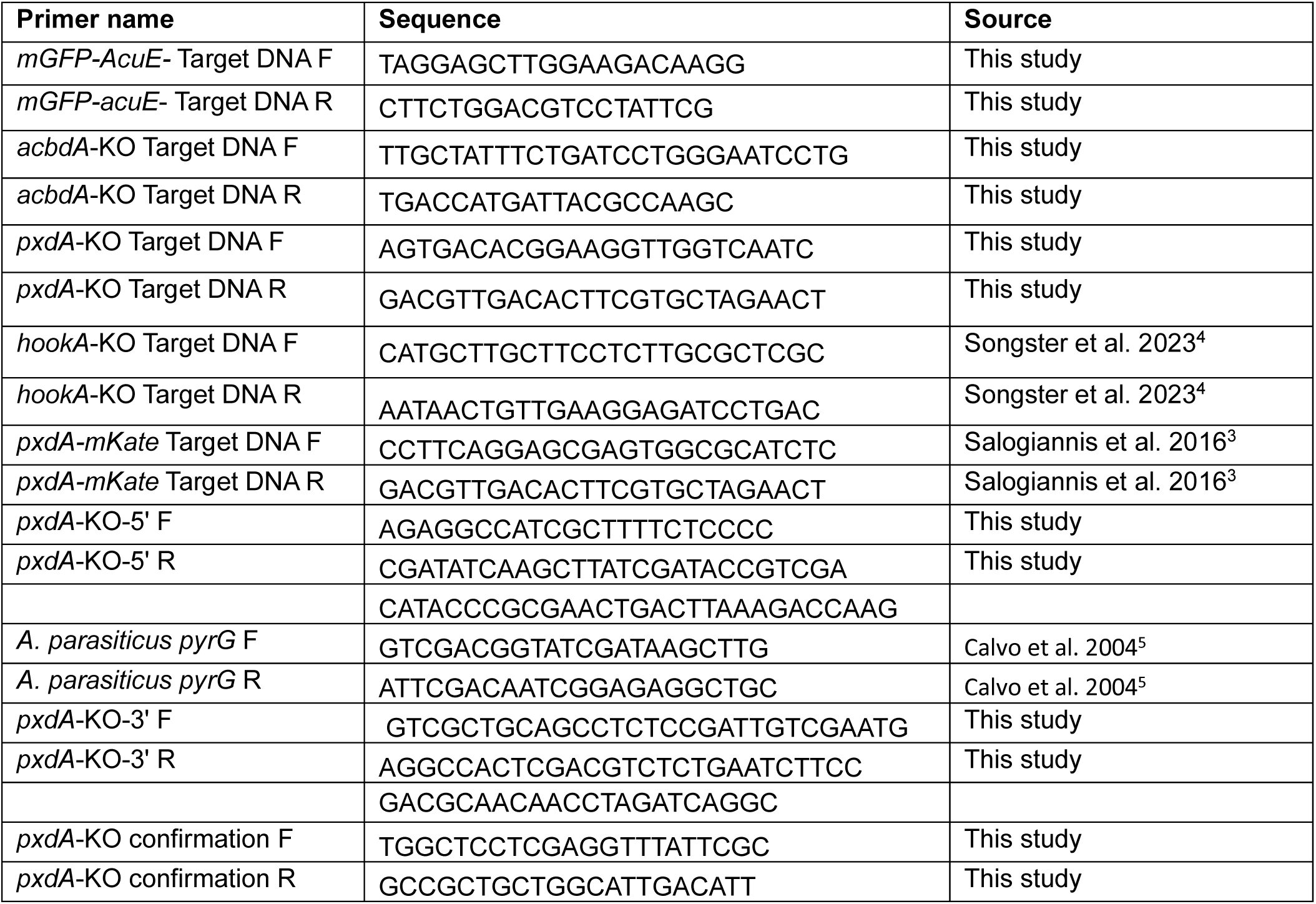
Primers used in this study.

